# Distinct temporal patterns of liver immune responses to pathogenic and non-pathogenic *Entamoeba histolytica* clones

**DOI:** 10.64898/2026.05.12.724513

**Authors:** Helena Fehling, Johannes Allweier, Barbara Honecker, Claudia Marggraff, Michel-Ruben Glagowski, Juliett Anders, Hannelore Lotter, Iris Bruchhaus

## Abstract

*Entamoeba histolytica* is a protozoan parasite that can cause severe liver disease known as amoebic liver abscess. However, only a subset of infected individuals develops invasive disease, indicating that host-parasite interactions are critical determinants of disease outcome. In this study, we investigated the clone-specific modulation of hepatic immune responses using non-pathogenic A1^np^ and pathogenic B2^p^ *E. histolytica* clones. Time-resolved transcriptome analyses (6, 12, 24 hours post-infection) in a murine model revealed distinct immune trajectories. Both clones activated innate immune pathways early after infection, but their responses differed markedly in magnitude and composition. A1^np^ infection induced a rapid and controlled inflammatory response associated with antimicrobial activity and resolution-promoting signalling. In contrast, B2^p^ infection triggered a stronger and more complex immune response characterised by pronounced cytokine and chemokine expression, activation of stress and redox pathways, and tissue remodelling processes. The B2^p^ induced response exhibited features of excessive immune activation, accompanied by the upregulation of counter-regulation genes such as *Ackr2*. These findings indicate that liver pathology is not solely determined by parasite presence, but rather may also be influenced by the nature and regulation of the host immune response. Overall, the observed differences between A1^np^ and B2^p^ infections suggest that parasite-specific properties shape hepatic immune activation and may influence disease progression.

**Author summary:** Although infection with the parasite *Entamoeba histolytica* can lead to severe liver disease, most infected individuals remain asymptomatic. This suggests that the outcome of the disease is not determined solely by the parasite, but also by how the host responds to the infection. In this study, we used a mouse model to compare how the liver reacts to infection with two *E. histolytica* clones that differ in their ability to cause amoebic liver abscesses. Using this model and time-resolved transcriptome analysis, we found that both clones trigger an early immune response; however, the nature of this response differs markedly. The non-pathogenic clone induced a rapid and controlled reaction associated with antimicrobial defence and tissue protection. In contrast, the pathogenic clone provoked a stronger and more prolonged inflammatory response accompanied by cellular stress and tissue remodelling processes. Notably, this heightened response also activated regulatory mechanisms that attempted to limit excessive inflammation. Our findings demonstrate that differences in disease severity are linked to the activation and regulation of the host immune system, rather than simply to the presence of the parasite.

## Introduction

*Entamoeba histolytica* is a protozoan parasite that causes intestinal and extraintestinal disease in humans. While most infections are asymptomatic, around 10% of affected individuals develop invasive manifestations, such as amoebic colitis or amoebic liver abscess (ALAs). The latter represents one of the most severe complications, accounting for substantial morbidity and mortality, particularly in endemic regions of Asia, Africa and Latin America, with an estimated 26,000 deaths worldwide in 2016 [1].

The life cycle of *E. histolytica* includes infectious cysts and motile trophozoites, which replicate within the intestinal lumen. Under conditions that are still poorly understood, the parasite can invade the intestinal mucosa and disseminate via the bloodstream to the liver. The pathogenesis of ALA is driven by a complex interplay between parasite-derived virulence factors and the host immune response.

Shortly after *E. histolytica* invades the liver, a pronounced infiltration of neutrophils and monocytes is observed [2-4]. These recruited cells, together with liver-resident Kupffer cells, play key roles in both early parasite control and tissue damage by rapidly inducing inflammatory mediators such as IL-6, IL-1β, TNF-α, IFN-γ, CXCL1 and CCL2, which promote the recruitment of additional immune cells and amplify the local inflammatory response [4-9]. Beyond their role in acute defence, dysregulated monocyte responses have been identified as critical drivers of hepatic immunopathology. In this context, Sellau *et al*. highlight that both the recruitment and functional polarisation of monocytes and neutrophils are key determinants of whether immune responses remain protective or shift towards tissue-damaging inflammation in the liver [10].

The activation of these innate immune cells is initiated through the recognition of amoebic components by pattern recognition receptors, including Toll-like receptors (TLRs), such as TLR2 and TLR4. Among the hepatic immune cell populations, Kupffer cells play a central role in early parasite sensing and the initiation of pro-inflammatory signalling [11, 12]. However, their activation can also exacerbate liver damage by promoting TNF production, thereby enhancing monocyte- and neutrophil-mediated tissue destruction. Consistently, depletion of Kupffer cells markedly reduces hepatic injury [2].

Neutrophils contribute to host defence against *E. histolytica* through the release of reactive oxygen species (ROS) and proteolytic enzymes, but these mechanisms can also promote collateral host tissue damage [13]. In addition, neutrophils can form neutrophil extracellular traps (NETs) in response to direct parasite contact or parasite-derived components. While NETs may immobilise amoebae, their effectiveness is limited, as *E. histolytica* expresses nucleases capable of degrading them [14-17]. Importantly, NET formation and neutrophil activation have been implicated in immunopathological processes, suggesting that excessive or dysregulated neutrophil responses may contribute to tissue injury rather than protection.

In addition to the host response, the virulence factors of *E. histolytica* play a decisive role in liver pathology. These include the Gal/GalNAc lectin, which mediates adhesion to host mucins in the gut and to host cells during tissue invasion, amoebic pore-forming proteins (amoebapores) that lyse target cells, and a 33-member family of papain-like cysteine peptidases (EhCPs) responsible for tissue degradation [18-22].

Experimental studies further suggest that different *E. histolytica* clones contribute to pathogenesis to varying degrees. Clonal lines derived from the reference strain HM-1:IMSS display substantial differences in virulence; some reliably induce liver abscesses in animal models, while others cause minimal or no lesions [18]. Transcriptomic comparison of the non-pathogenic clone A1^np^ and the pathogenic clone B2^p^ identified 76 differentially expressed genes (46 upregulated in A1^np^ versus B2^p^ and 30 upregulated in B2^p^ versus A1^np^) [18]. Functional assays using overexpression transfectants identified eight proteins that modulate pathogenicity, including a metallopeptidase, C2 domain containing proteins, alcohol dehydrogenases, and several hypothetical proteins [18]. Furthermore, RNA interference approaches were used to analyze the contribution of genes differentially expressed between A1^np^ and B2^p^ trophozoites to pathogenicity. Trigger-induced RNAi-mediated gene silencing of 15 candidates revealed specific effects on ALA formation in a mouse model [23].

However, it remains unclear whether and how the hepatic host response differs in a clone-specific manner between infections with pathogenic and non-pathogenic *E. histolytica* trophozoites. While differences in parasite virulence and selected host responses have been described, a systematic and time-resolved comparison of the global host response, particularly at the transcriptional level, is still lacking.

This study aimed to analyse the temporal dynamics of the hepatic response to infection with non-pathogenic (A1^np^) and pathogenic (B2^p^) *E. histolytica* trophozoites using a murine model. By performing transcriptomic analyses at three time points (6, 12 and 24 hours post infection (hpi)) after intrahepatic infection, we sought to determine how parasite-specific differences shape the hepatic immune profile. The data revealed shared patterns of innate immune activation as well as clone-specific differences in the magnitude, composition and temporal dynamics, highlighting clone-specific modulation of host immunity. While A1^np^ trophozoites triggered a rapid, classical inflammatory response with a strong antimicrobial component, B2^p^ induced a markedly enhanced and complex immune reaction characterised by a combination of pro-inflammatory signalling, cellular stress pathways, and counter-regulatory mechanisms. These findings emphasise that distinct *E. histolytica* clones elicit unique host immune responses, which play a crucial role in determining disease progression and the extent of liver tissue damage.

## Methods

### *E. histolytica* cell culture

*E. histolytica* trophozoites were maintained axenically under microaerophilic conditions at 37°C in TYI-S-33 medium [24] using plastic culture flasks (Corning, Kaiserslautern, Germany). Non-pathogenic clone A1^np^ and pathogenic clone B2^p^ originated from the HM-1:IMSS-A and HM-1:IMSS-B cell lines, respectively [18].

### ALA formation in mice

Animal infections were performed with 10-to 12-week-old male C57BL/6 mice bred in the animal facility of the Bernhard Nocht Institute for Tropical Medicine (Hamburg, Germany). All procedures were approved by the institutional review board of the State of Hamburg, Germany (Ministry of Health and Consumer Protection; approval no. 145/13, issued 20 January 2014) and conducted in accordance with institutional regulations and the ARRIVE guidelines (https://www.nc3rs.org.uk/arrive-guidelines).

For infection experiments, 1 × 10^6^ *E. histolytica* trophozoites were cultured in 75 ml culture flasks for 24 hours. After incubation, the cells were harvested and resuspended in incomplete TYI-S-33 medium. Then, 1.25 × 10^5^ trophozoites in 25 μl of incomplete TYI-S-33 medium were injected directly into the left liver lobe as previously described [4, 18]. The mice were sacrificed at 6, 12 and 24 hpi, after which the infected liver lobe was transferred either into TRIzol reagent (Thermo Fisher Scientific) for RNA extraction or fixed in 4% formalin for immunohistochemistry.

Since no sham-injected control group was included, all control comparisons were made against untreated animals. To account for potential procedure-related effects, particular emphasis was placed on the direct comparison between A1^np^ and B2^p^ infections.

### RNA extraction and RNAseq

The liver tissue was homogenised in 500 μL of TRIzol reagent (Thermo Fisher Scientific). An additional 500 μL of TRIzol was added after homogenisation. RNA was then isolated using the mirVana™ miRNA Isolation Kit (Ambion), following the manufacturer’s protocol. RNA concentration and integrity were assessed spectrophotometrically using a NanoDrop 2000 (Thermo Fisher Scientific) and with an Agilent 2100 Bioanalyzer employing the RNA 6000 Pico Assay kit (Agilent Technologies). Potential genomic DNA contamination was eliminated using the TURBO DNA-free Kit (Thermo Fisher Scientific, Hampton, NH, USA). Library preparation and high-throughput RNA sequencing were conducted by BGI (Shenzhen, China) using the Illumina HiSeq 4000 platform with paired-end 100 bp reads (PE100).

### Bioinformatic Analyses/Statistics

RNA-seq data was processed using CLC Genomics Workbench v24.0 (QIAGEN). Initially, raw sequencing reads were adapter-trimmed and quality-filtered with a quality score of 0.05. Reads were subsequently mapped to the *Mus musculus* reference genome (GRCm39), and gene expression was quantified in Transcripts Per Million (TPM). Heatmaps were created using the R/Shiny-based tool Heatmapper [25]. Only those genes with a differential expression with a fold change of ≥ 2.0, a FDR-adjusted *p* value of ≤ 0.05 and a minimum TPM threshold of ≥ 2.0 were included in this analysis. The interaction network analysis was performed with the database STRING, version 12.0 [26]. The following setting was used: Active interaction sources: text mining, experiments, databases, co-expression; Meaning of network edges: high confidence 0.700; Network display mode: Interactive; Network display options: Hide disconnected nodes in the network; Clustering Options: MCL clustering, inflation parameter 3. Visualization of functional enrichment for Reactome and KEGG pathways was performed using STRING. Volcano plots for the visualization of significantly differentially expressed genes were created using the European Galaxy Server (https://rna.usegalaxy.eu/) [27].

### Immunohistology

Liver tissue from mice infected with pathogenic B2^p^ trophozoites was fixed in formalin (4%) and embedded in paraffin. Sections (0.2 mm) were stained with H&E (Hematoxylin and Eosin), or prepared for immunohistochemistry. Immunohistochemistry was performed using the following primary antibodies: anti-Ackr2 (1:400, Proteintech), anti-Rgs16 (1:400, Biorbyt), anti-Btg2 (1:400, Biorbyt), anti-Gadd45b (1:200, MyBioSource) and anti-Sesn1 (1:400, Biorbyt). Detection was carried out with the Zytomed Chem-Plus HRP Polymer Kit (Zytomed) following the manufacturer’s instructions, and visualisation was achieved using 3,3′-diaminobenzidine substrate solution (Dako).

## Results

### Strong and clone-dependent transcriptional responses in the liver following infection with *E. histolytica* trophozoites

In order to investigate whether pathogenic and non-pathogenic amoebae elicit distinct hepatic responses, male mice aged 10 to 12 weeks were injected intrahepatically with 1.25×10^5^ *E. histolytica* trophozoites of either the non-pathogenic A1^np^ clone or the pathogenic B2^p^ clone. Liver tissue was collected at 6, 12 and 24 hpi and subjected to transcriptomic analysis. The primary aim was to define clone-specific differences in the hepatic host response by directly comparing A1^np^ and B2^p^ infections across time. In addition, differentially expressed genes were identified relative to non-infected control mice, using a fold change (FC) threshold of ≥2.0 and a false discovery rate (FDR) threshold of < 0.05.

A marked hepatic response was detectable as early as 6 hpi. Following the injection of non-pathogenic A1^np^ trophozoites, 687 genes were found to be upregulated and 814 downregulated in comparison to the control group. In contrast, infection with pathogenic B2^p^ trophozoites resulted in 814 genes being upregulated and 952 being downregulated. At 12 hpi, the total number of differentially expressed genes increased from 1,501 to 1,689 in the A1^np^ group and from 1,766 to 2,643 in the B2^p^ group. After 24 hpi, the number of differentially expressed genes decreased considerably. In the A1^np^ group, 215 genes were upregulated and 127 were downregulated, making a total of 342 genes. In the B2^p^ group, 296 genes were upregulated and 236 were found to be downregulated, making a total of 532 genes (Fig 1; S1 Table). Direct comparison of differential gene expression between A1^np^ and B2^p^ revealed only 12 genes at 6 hpi, but this number increased markedly to 543 genes at 12 hpi and further to 836 genes at 24 hpi (Fig 1; S1 Table).

**Fig. 1.**
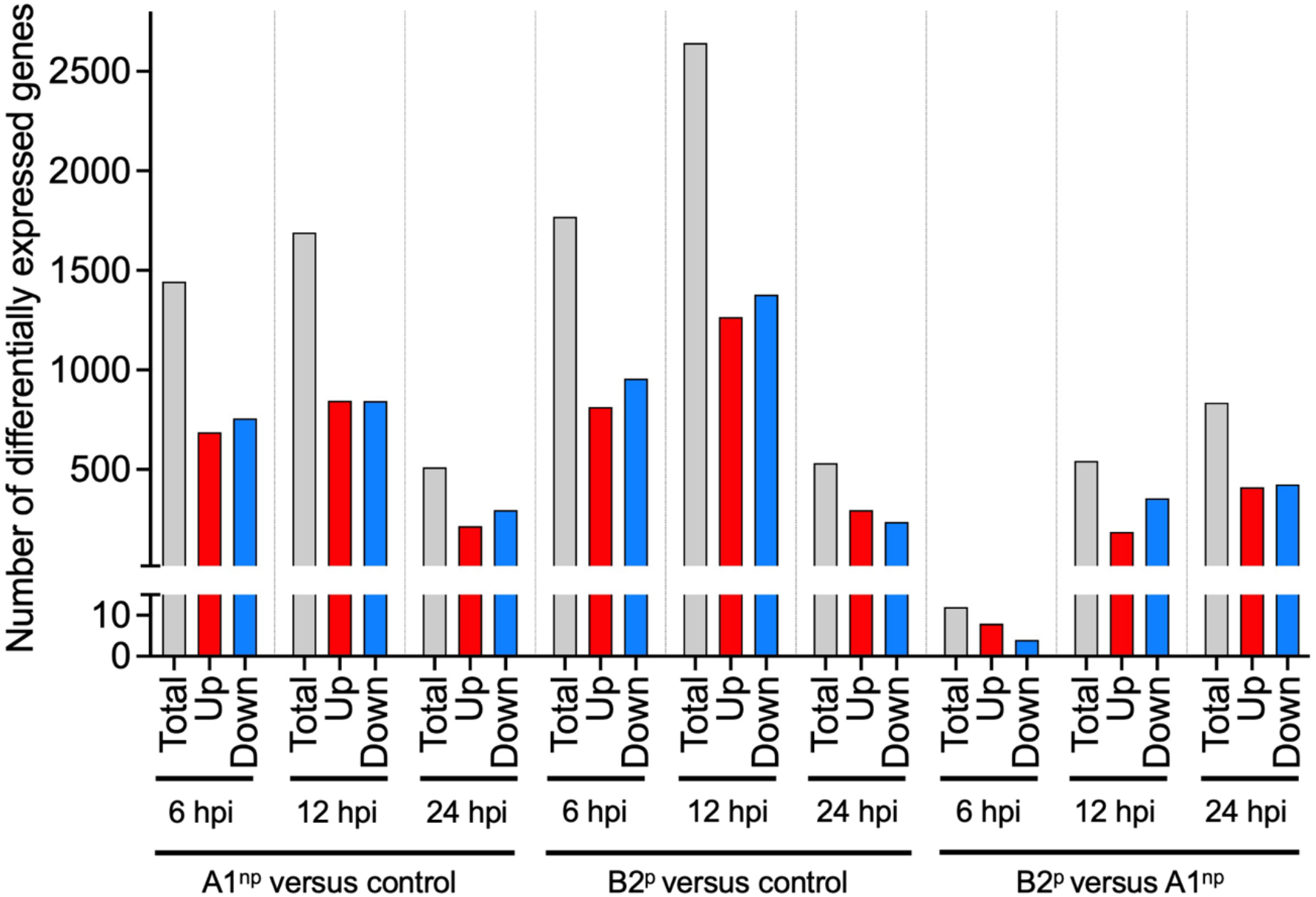
Differential gene expression in liver tissue following infection with A1^np^ and B2^p^ trophozoites. The number of differentially expressed genes (DEGs) in liver tissue of mice intrahepatically infected with *Entamoeba histolytica* clones A1^np^ and B2^p^ is shown at 6, 12, and 24 hours post-infection (hpi). Differential expression analysis was performed relative to uninfected control mice (A1^np^ versus control; B2^p^ versus control) and between infections (B2^p^ versus A1^np^). Bars represent the total number of DEGs (grey), as well as upregulated (red) and downregulated (blue) genes. Differential expression was defined as ≥2-fold change with FDR < 0.05 (control: n = 5; infected groups: n = 4 per time point).

Reactome and KEGG enrichment pathway analyses of significantly upregulated genes revealed a rapid and pronounced hepatic response to *E. histolytica* infection already at 6 hpi (Fig 2A/B). In both A1^np^ and B2^p^ infections, KEGG analysis revealed enrichment of central signalling pathways, including MAPK, TNF, NF-κB, FoxO, PPAR and IL-17 signalling, indicating activation of core inflammatory and regulatory networks (Fig 2A). Consistently, Reactome analysis highlighted shared enrichment of pathways related to Toll-like receptor (TLR) signalling, cytokine and interleukin signalling, as well as platelet activation and degranulation, alongside metabolic processes (Fig 2B), reflecting a combined inflammatory and metabolic host response.

**Fig. 2.**
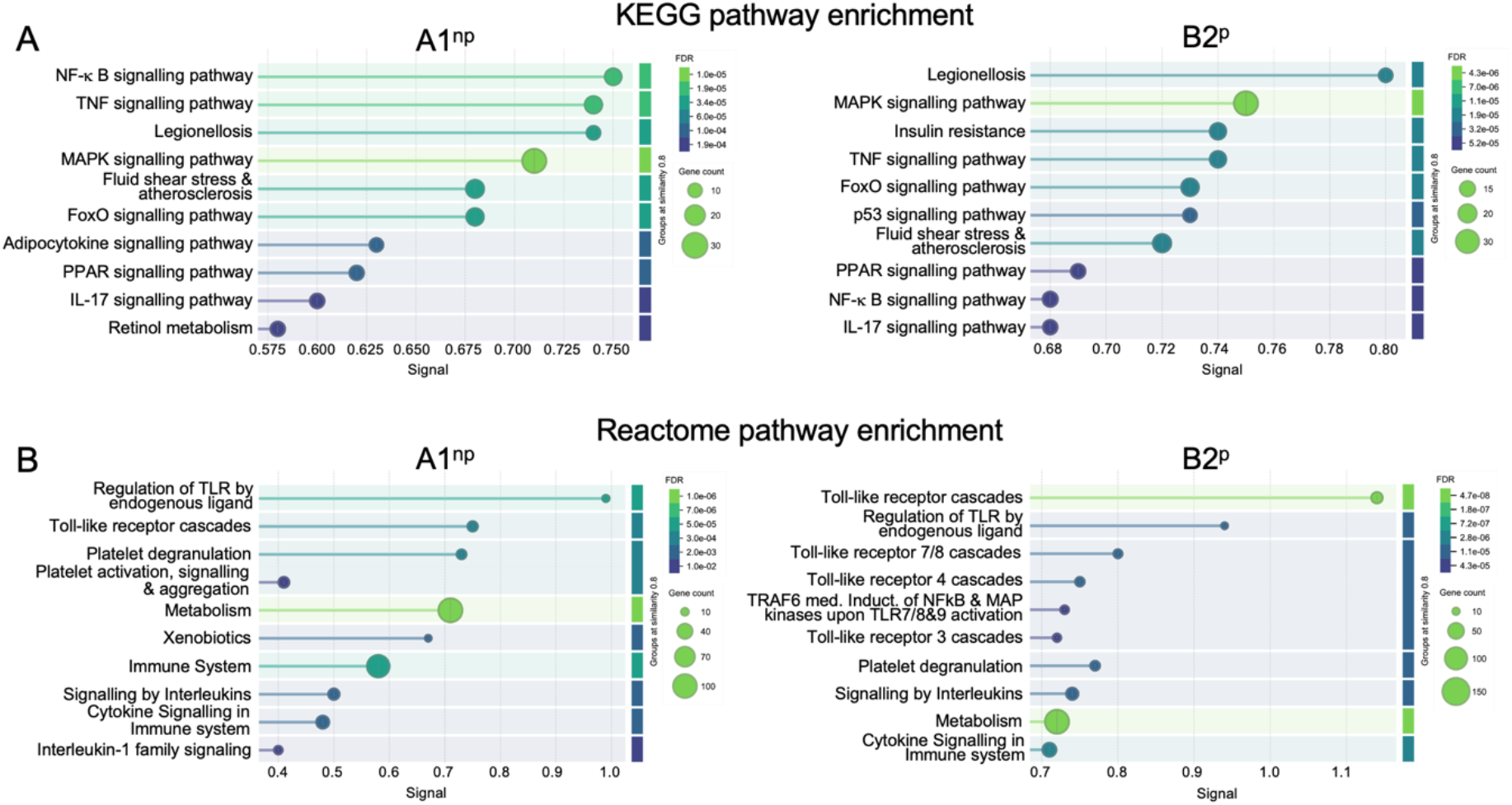
KEGG and Reactome pathway enrichment analysis of genes upregulated at 6 hpi in mice infected with non-pathogenic (A1^np^) and pathogenic (B2^p^) *E. histolytica* trophozoites compared to uninfected controls. KEGG (A) and Reactome (B) pathway enrichment analysis was performed using all genes that were significantly upregulated at 6 hpi in mouse liver tissue in comparison to the uninfected control (fold change (FC) ≥ 2.0; false discovery rate (FDR) < 0.05). The pathways are shown for the non-pathogenic clone A1^np^ (left panels) and the pathogenic clone B2^p^ (right panels). The x-axis represents the enrichment signal, reflecting the strength of pathway overrepresentation. Dot size indicates the number of genes associated with each pathway (gene count), and the colour gradient represents the FDR-adjusted *p*-value. Control: n = 5; A1^np^ and B2^p^: n = 4.

However, differences in the hepatic response to A1^np^ and B2^p^ trophozoite infection are already detectable at 6 hpi. In A1^np^-infected livers, KEGG analyses show a comparatively stronger representation of pathways related to retinol metabolism and adipocytokine signalling (Fig 2A/B). This pattern may suggest a more regulated, metabolically integrated host response associated with immune modulation and maintenance of tissue homeostasis. Although cytokine- and interleukin-associated pathways are also enriched, they appear less prominent relative to metabolic pathways. In contrast, B2^p^ infection is characterized by a comparatively stronger enrichment of immune-related pathways, including multiple Toll-like receptor cascades, together with insulin and p53 signalling pathways (Fig 2A/B), indicating a shift in the relative representation of immune-associated processes.

At 12 hpi, the hepatic transcriptional responses to A1^np^ and B2^p^ trophozoites diverge increasingly. Only three KEGG pathways remain enriched in both groups: IL-17 signalling, AGE-RAGE signalling in diabetic complications, and TNF signalling pathways (Fig 3A). Conversely, Reactome analysis revealed shared Toll-like receptor cascades, and pathways of the immune system, indicating conserved engagement of core innate immune and pattern recognition mechanisms despite differing infection profiles (Fig 3B).

**Fig. 3.**
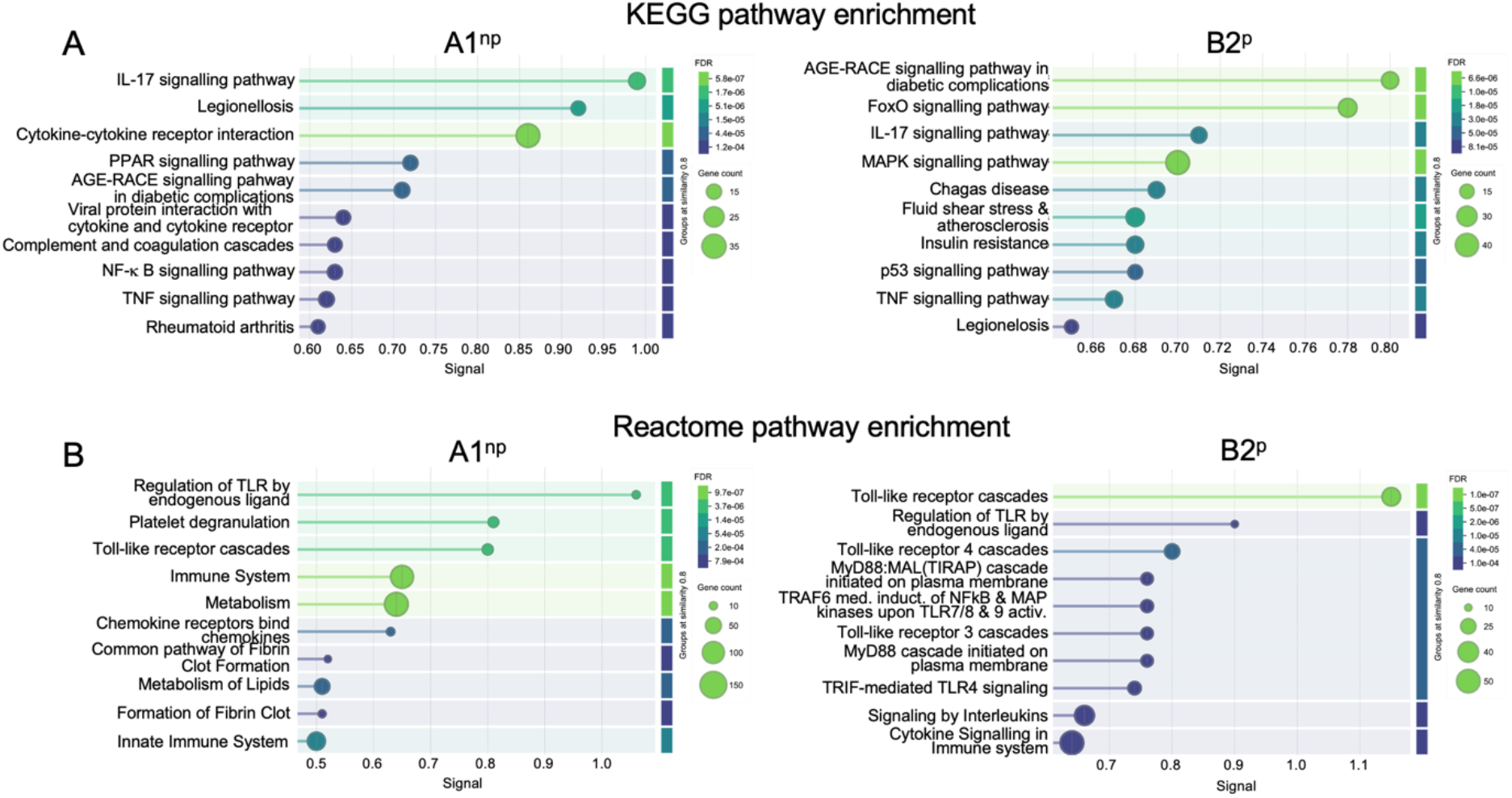
KEGG and Reactome pathway enrichment analysis of genes upregulated at 12 hpi in mice infected with non-pathogenic (A1^np^) and pathogenic (B2^p^) *E. histolytica* trophozoites compared to uninfected controls. KEGG (A) and Reactome (B) pathway enrichment analysis was performed using all genes that were significantly upregulated at 12 hpi in mouse liver tissue in comparison to the uninfected control (FC ≥ 2.0; FDR ≤ 0.05). The pathways are presented for the non-pathogenic clone A1^np^ (left panels) and the pathogenic clone B2^p^ (right panels). The x-axis represents the enrichment signal, reflecting the strength of pathway overrepresentation. Dot size indicates the number of genes associated with each pathway (gene count), and the colour gradient represents the adjusted *p*-value (FDR). Control: n = 5; A1^np^ and B2^p^: n = 4.

However, the response to the two *E. histolytica* clones differs in a number of ways. In response to A1^np^ infection, enriched KEGG pathways include cytokine-cytokine receptor interaction PPAR signalling pathway, viral protein interaction with cytokine and cytokine receptor, complement and coagulation cascades, NF-κB signalling pathway and rheumatoid arthritis (Fig 3A). Reactome pathways reflect a similar immunovascular activation profile, with enrichment of platelet degranulation, pathways of fibrin clot formation, metabolism of lipids and metabolism (Fig 3B). These findings suggest a coordinated vascular-inflammatory response, marked by chemotactic signalling, coagulation involvement, and lipid-mediated immune modulation.

By contrast, B2^p^ infection predominantly activates KEGG pathways associated with cellular stress and regulatory control, including the FoxO, MAPK and p53 signalling pathways (Fig 3A). Reactome analysis further revealed the targeted activation of specific Toll-like receptors, including TLR3, TLR4, and downstream TRAF6- and the MyD88/MAL-dependent signalling axes (Fig 3B). Together, these responses indicate a state of pronounced cellular dysregulation characterised by stress signalling, cell cycle arrest, metabolic disturbance, DNA damage and apoptosis, accompanied by robust, TLR-driven pro-inflammatory cascades.

At 24 hpi, liver cells continued to exhibit distinct transcriptional responses to infection with A1^np^ and B2^p^ trophozoites. While some overlap was observed, including the enrichment of KEGG pathways such as the IL-17 signalling pathway, viral protein interaction with cytokine and cytokine receptors, retinol metabolism, and legionellosis, clear differences between the two infection conditions remained evident (Fig 4A).

**Fig. 4.**
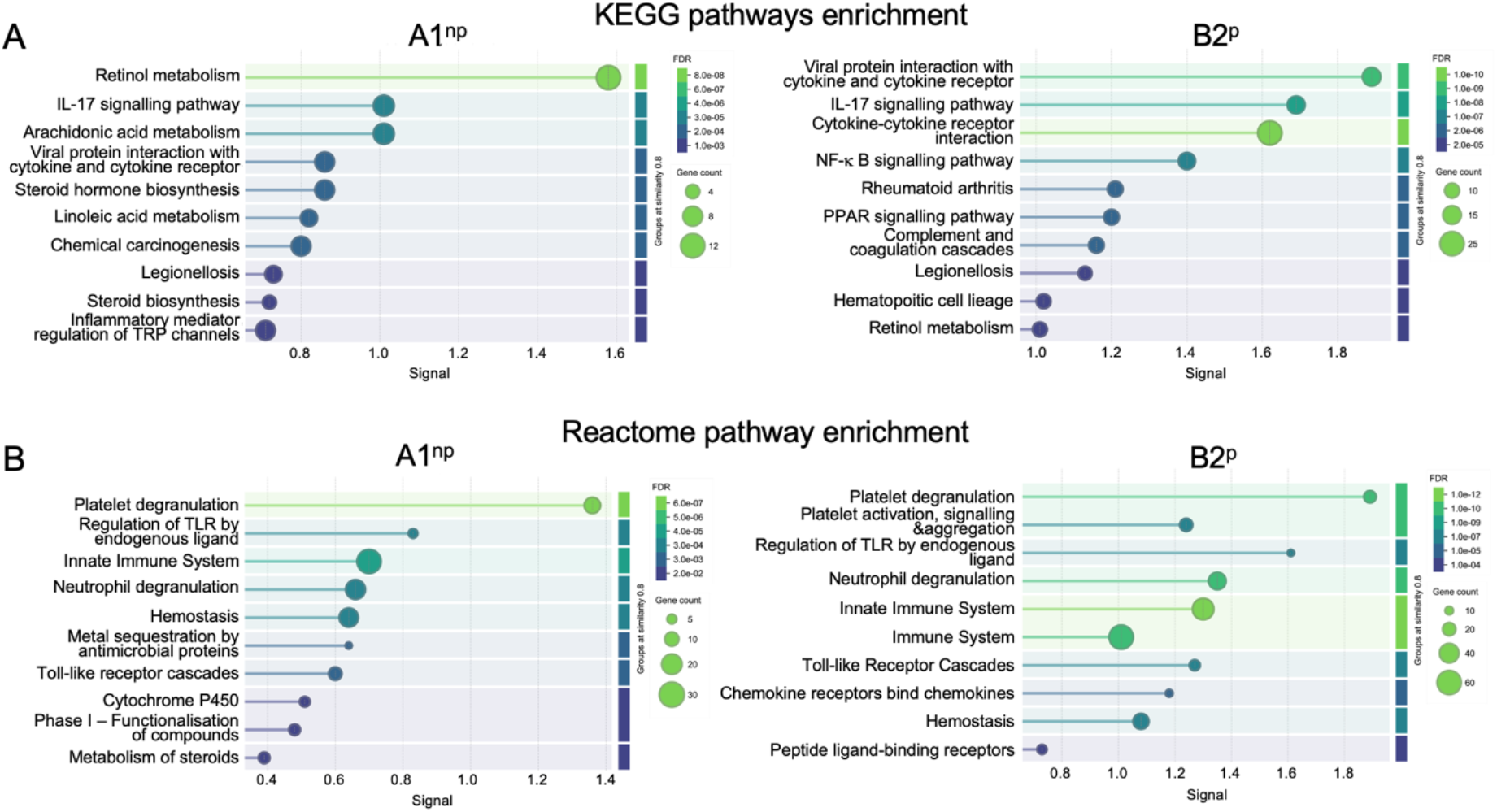
KEGG and Reactome pathway enrichment analysis of genes upregulated at 24 hpi in mice infected with non-pathogenic (A1^np^) and pathogenic (B2^p^) *E. histolytica* trophozoites compared to uninfected controls. KEGG (A) and Reactome (B) pathway enrichment analysis was performed using all genes that were significantly upregulated at 24 hpi in mouse liver tissue in comparison to the uninfected control (FC ≥ 2.0; FDR ≤ 0.05). The pathways are presented for the non-pathogenic clone A1^np^ (left panels) and the pathogenic clone B2^p^ (right panels). The x-axis represents the enrichment signal, reflecting the strength of pathway overrepresentation. Dot size indicates the number of genes associated with each pathway (gene count), and the colour gradient represents the adjusted *p*-value (FDR). Control: n = 5; A1^np^ and B2^p^: n = 4.

Infection with A1^np^ trophozoites was characterised by the specific enrichment of metabolic pathways, including arachidonic acid metabolism, steroid hormone biosynthesis, and linoleic acid metabolism. In contrast, B2^p^ infection predominantly induced immune-related KEGG pathways, such as cytokine–cytokine receptor interaction, NF-κB signalling, rheumatoid arthritis, PPAR signalling, complement and coagulation cascades, and hematopoietic cell lineage (Fig 4A). According to the Reactome analysis, a set of co-regulated pathways was identified, including platelet degranulation, regulation of Toll-like receptor (TLR) signalling by endogenous ligands, the immune system, neutrophil degranulation, and hemostasis. Pathways specifically enriched in A1^np^ infection were primarily related to metabolic and detoxification processes, including metal sequestration by antimicrobial proteins, cytochrome P450, phase I functionalization of compounds, and steroid metabolism (Fig 4B). In contrast, B2^p^ infection led to the additional enrichment of pathways associated with immune cell communication and signalling, such as chemokine receptors, chemokine-binding receptors, and peptide ligand-binding receptors (Fig 4B).

### Clone-specific gene expression reveals divergent immune activation in response to A1^np^ and B2^p^ infection

Taken together, these pathway-based analyses comparing infected livers to untreated controls highlight distinct immune-related responses induced by A1^np^ and B2^p^ trophozoites. A1^np^ is associated with regulatory and protective pathways, whereas B2^p^ elicits a comparatively stronger pro-inflammatory activation. To further explore the molecular basis of these responses, we examined immune-related gene expression. Of the thousands of genes differentially expressed in liver cells following infection with A1^np^ and B2^p^ trophozoites, 151 could be linked to immune pathways (Table S3), the majority of which were upregulated in both infection groups.

Although distinct expression patterns also emerged over time, several genes were similarly regulated in response to both A1^np^ and B2^p^ trophozoites over the course of infection. At 6 hpi, shared transcriptional responses included the significant upregulation of genes involved in sensing and regulating the innate immune response. These genes included *Clec4e, Cd14, Irak3, Cd33*, and *Cd300lf* (Fig 5A, S3 Table). Several inflammation- and metabolism-associated genes were also co-regulated, including *Cxcl2, Acod1, Serpina3g/n/m* and *Slfn4*, all of which showed similarly high fold changes in both groups (Fig 5A-C, S3 Table).

**Fig. 5.**
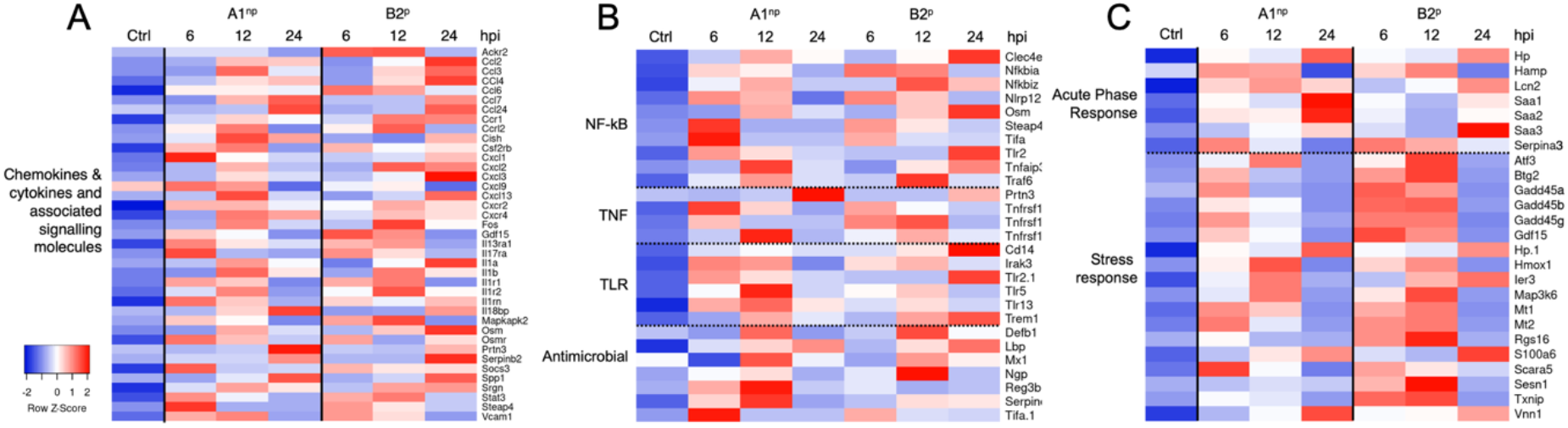
Time-resolved hepatic gene expression responses to infection with A1^np^ and B2^p^ trophozoites. (A-C) Heatmaps showing the average expression of selected genes in liver tissue at 6, 12, and 24 hpi infected either with non-pathogenic (A1^p^) or pathogenic (B2^p^) *E. histolytica* trophozoites (n = 4). Expression values are shown as Z-scores for each row, calculated from normalised TPMs (transcripts per million). Red indicates high gene expression and blue indicates low gene expression (ctrl; uninfected mice, n = 5). Genes are grouped according to functional categories: (A) chemokines, cytokines, and associated signalling molecules; (B) key inflammatory pathways including NF-κB, TNF, TLR signalling, and antimicrobial responses; (C) acute phase and stress response genes.

At 12 hpi, a subset of these genes remained commonly upregulated, such as *Il1rn, Cxcl1, Cxcr4, C5ar1, Csf3r*, and again *Serpina3m*, indicating a sustained inflammatory and regulatory response in both infection settings (Fig 5A/B; S3 Table).

Further overlap in gene regulation was observed between the two groups by 24 hpi, including for *Cxcr4, Csf3r, Hp, Itgam, Cish, Cd33*, and *Gas6* (Fig 5A/B, S3 Table).

Despite these similarities, clear differences in the dynamics and overall magnitude of gene expression profiles between A1^np^ and B2^p^ infections were evident at most time points (Fig 1). This highlights the fundamentally different nature of the liver’s response to non-pathogenic and pathogenic *E. histolytica* clones.

As early as 6 hpi, distinct differences in gene expression emerged between the two infection groups, with several genes being more strongly upregulated following B2^p^ infection. These included *Il1r2, Rgs16, Cxcl3, Gadd45b, Gadd45g, Txnip, Mmp8, Osm* and *Igfbp1*. In contrast, stronger expression in the A1^np^ group was observed for *Serpine1, Lcn2*, and *Cxcl1* (Fig 5A, S3 Table).

At 12 hpi, the expression of genes encoding chemokines and their receptors was again more pronounced in the B2^p^ group, including *Cxcl2, Ccr1*, and *Cxcr2*. In contrast, *Ccl7* and *Cxcl3* showed slightly higher expression in the A1^np^ group (Fig 5A, S3 Table). *Ackr2*, which encodes an atypical chemokine receptor involved in chemokine degradation, was also induced more strongly following B2^p^ infection (Fig 5A, S3 Table). Genes involved in interleukin signalling exhibited a consistent trend towards higher expression in the B2^p^ group, including *Il1b, Il1r1, Il17ra*, and *Il1r2* while *Il1rn* was similarly expressed in both groups (Fig 5A, S3 Table). The expression of genes involved in signal transduction pathways was also more strongly induced by B2^p^ infection. These genes included *Nfkbia, Ripk, Traf6*, and *Socs2* (Fig 5B, S3 Table). Additional genes associated with T cell regulation and inflammation, such as *Tnfrsf1b, Phf11*, and *Rgs16*, were also more strongly expressed in the B2^p^ group (Fig 5B/C, S3 Table). Similarly, genes associated with the stress response and redox including *Gadd45a, Gadd45b, Gadd45g, Txnip, Mt1*, and *Mt2* were also upregulated to a greater extent in the B2^p^ group (Fig 5C, S3 Table). In terms of antimicrobial responses, *Reg3b* showed a significant increase following A1^np^ infection, whereas *Ngp*, a neutrophil-specific antimicrobial protein, exhibited stronger induction in the B2^p^ group (Fig 5B, S3 Table).

This pattern of heightened gene expression in the B2^p^ group persisted at 24 hpi. There was stronger induction for *Cxcl3, Cxcl2, Ccl3, Osm, Il1r2*, and *Trem1*. Additionally, pro-inflammatory and innate immunity-related genes, including *Ptgs2, Saa3, Clec4d, Clec4e*, and *Serpine1* exhibited higher expression following B2^p^ infection. An exception to this trend was *Saa2*, which exhibited higher expression in the A1^np^ group (Fig 5A-C; S3 Table).

### Temporal progression of inflammatory networks

Fig 6 shows the temporal development of gene expression (fold change compared to control) of a STRING cluster containing genes associated with neutrophil chemotaxis, the interaction of viral proteins with cytokines and cytokine receptors, and the inflammatory response. To simplify the presentation, the absolute expression levels are not shown. Instead, Fig 6 shows whether a gene is similarly regulated in both infections (grey), is more strongly upregulated after A1^np^ infection (light blue), or after B2^p^ infection (light green). Genes that are expressed exclusively after infection with one clone are marked in dark blue (A1^np^) or dark green (B2^p^). The regulatory dynamics described above are again evident. After 6 hpi, 23 genes show comparable regulation in both infection groups, while seven genes in the B2^p^ group show higher expression than in the A1^np^ group and five genes are more strongly expressed after infection with A1^np^ than after infection with B2^p^. At 12 hpi, 27 genes remain similarly regulated, but 15 genes now show increased expression in the B2^p^ group, compared to only five in the A1^np^ group. At 24 hpi, the number of genes with similar regulation drops to 12, while 24 genes are more highly expressed after B2^p^ infection. At this point, no gene shows increased expression specifically in the A1^np^ group compared to the B2^p^ group.

**Fig. 6.**
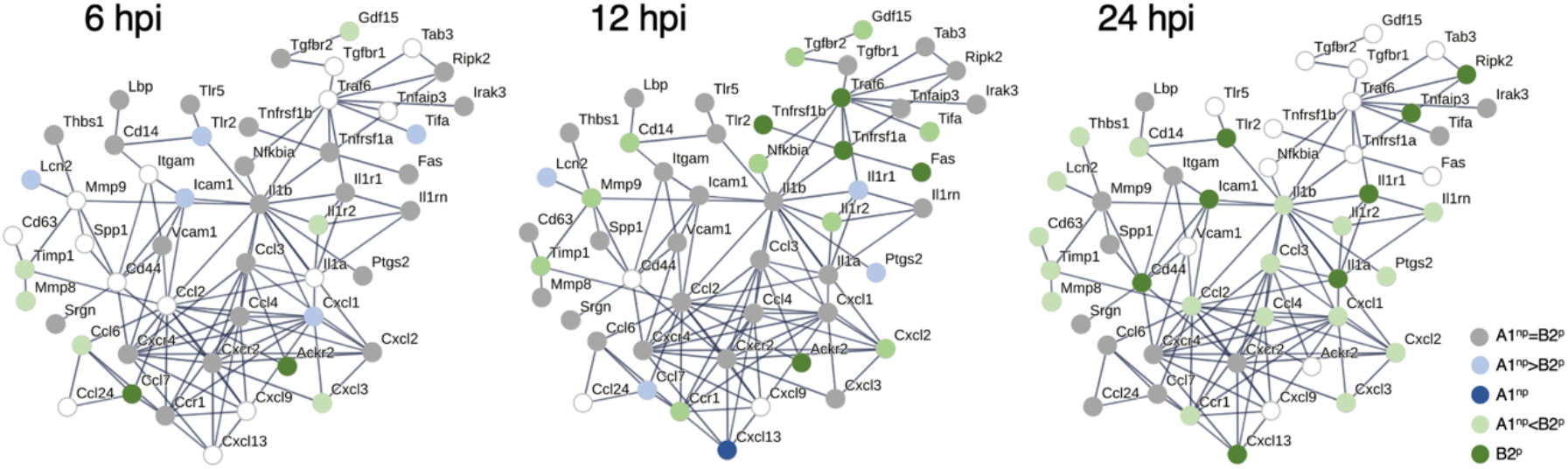
STRING cluster of neutrophil chemotaxis and cytokine signalling. The STRING cluster comprises genes associated with neutrophil chemotaxis, cytokine signalling and inflammatory responses. Temporal regulation at 6, 12 and 24 hpi is shown. Genes that do not exhibit differential expression during infection with A1^np^ or B2^p^ trophozoites in comparison to the control are represented as unfilled circles. Genes with similar differential expression after A1^np^ and B2^p^ infection are shown as grey circles (FC < 2.0). Those more strongly expressed after A1^np^ infection than B2^p^ infection are shown in light blue. Genes that are more strongly expressed after B2^p^ infection than after A1^np^ infection are shown in light green Genes that are exclusively expressed following infection by one clone only are marked in dark blue for A1^np^ trophozoites and in dark green for B2^p^ trophozoites.

### Time-dependent differences in biological processes between A1^np^ and B2^p^ infections

However, a subset of genes was not only more strongly expressed in B2^p^-infected than in A1^np^-infected mice, but also remained differentially expressed in the direct comparison between the two clones. At 6 hpi, only eight genes were upregulated in liver cells following B2^p^ infection compared to A1^np^, whereas four genes were downregulated. By 12 hpi, this number had increased markedly, with 187 genes being upregulated and 356 genes being downregulated in B2^p^-infected mice compared to A1^np^-infected mice. At 24 hpi, the numbers of up- and downregulated genes converged, with 411 genes being upregulated and 425 being downregulated (Fig 1, Fig 7A-C, S1 Table).

**Fig. 7.**
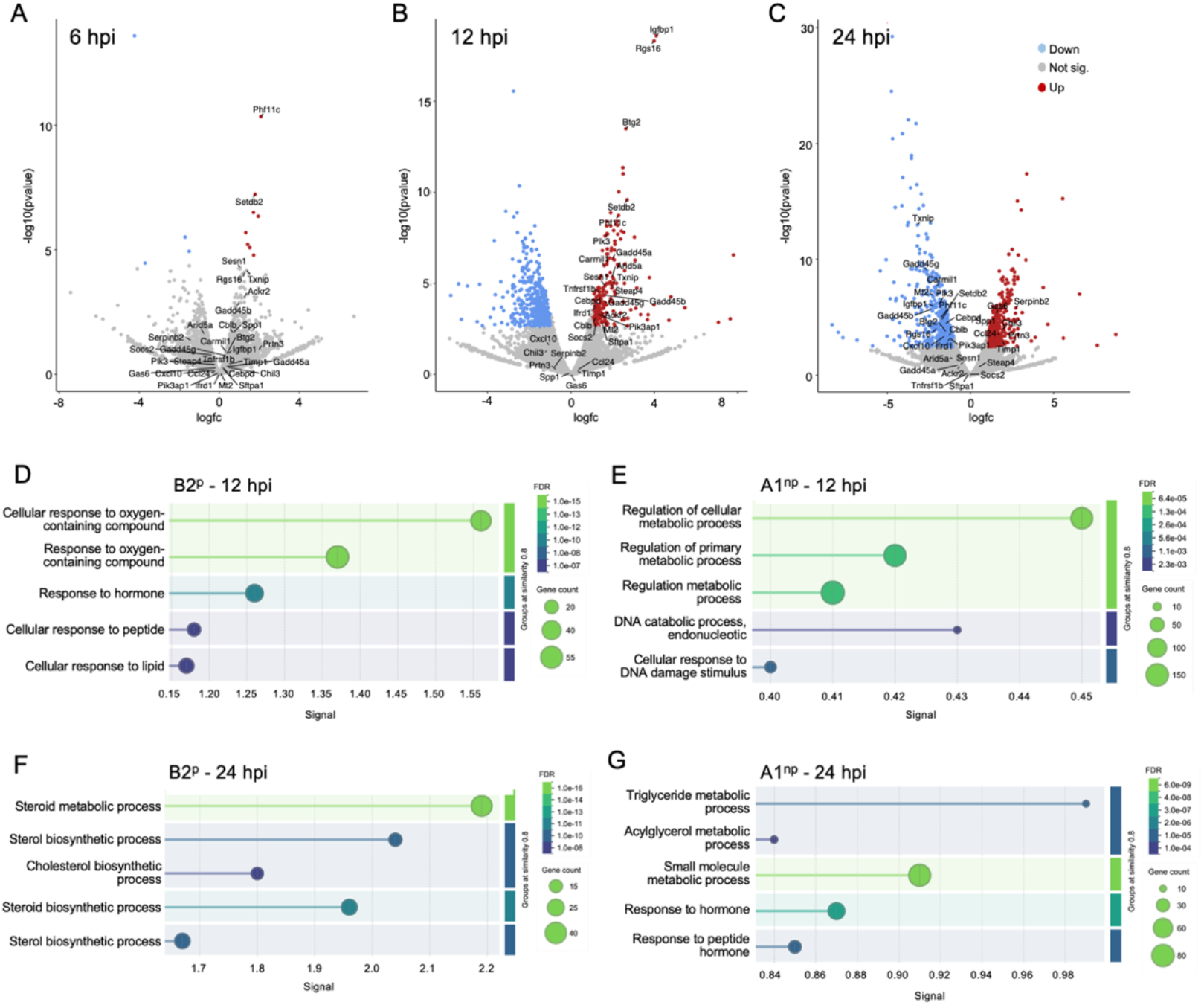
Differential gene expression and enrichment of biological processes in liver tissue following infection with A1^np^ and B2^p^ trophozoites. (A–C) Volcano plots showing differentially expressed genes (FC ≥ 2, FDR < 0.05) in liver cells at 6 (A), 12 (B) and 24 (C) hpi in the direct comparison between B2^p^- and A1^np^-infected mice. Each dot represents one gene, plotted according to log fold change (logFC) versus –log_10_(p-value). Significantly upregulated genes in B2^p^ compared to A1^np^ are shown in red, downregulated genes in blue and non-significantly regulated genes in grey. (D–G) Gene ontology (GO) term biological processes enriched in genes that were differentially expressed between A1^np^-and B2^p^-infected mice at 12 and 24 hpi. The left panels (D, F) show significantly enriched GO biological process terms for genes that are upregulated in B2^p^-infected mice relative to A1^np^-infected mice, and the right panels show the same for genes that are upregulated in A1^np^-infected mice relative to B2^p^-infected mice (E, G). The data are presented separately for 12 hpi (upper panels, D, E) and 24 hpi (lower panels, F, G).

**Figure 8.**
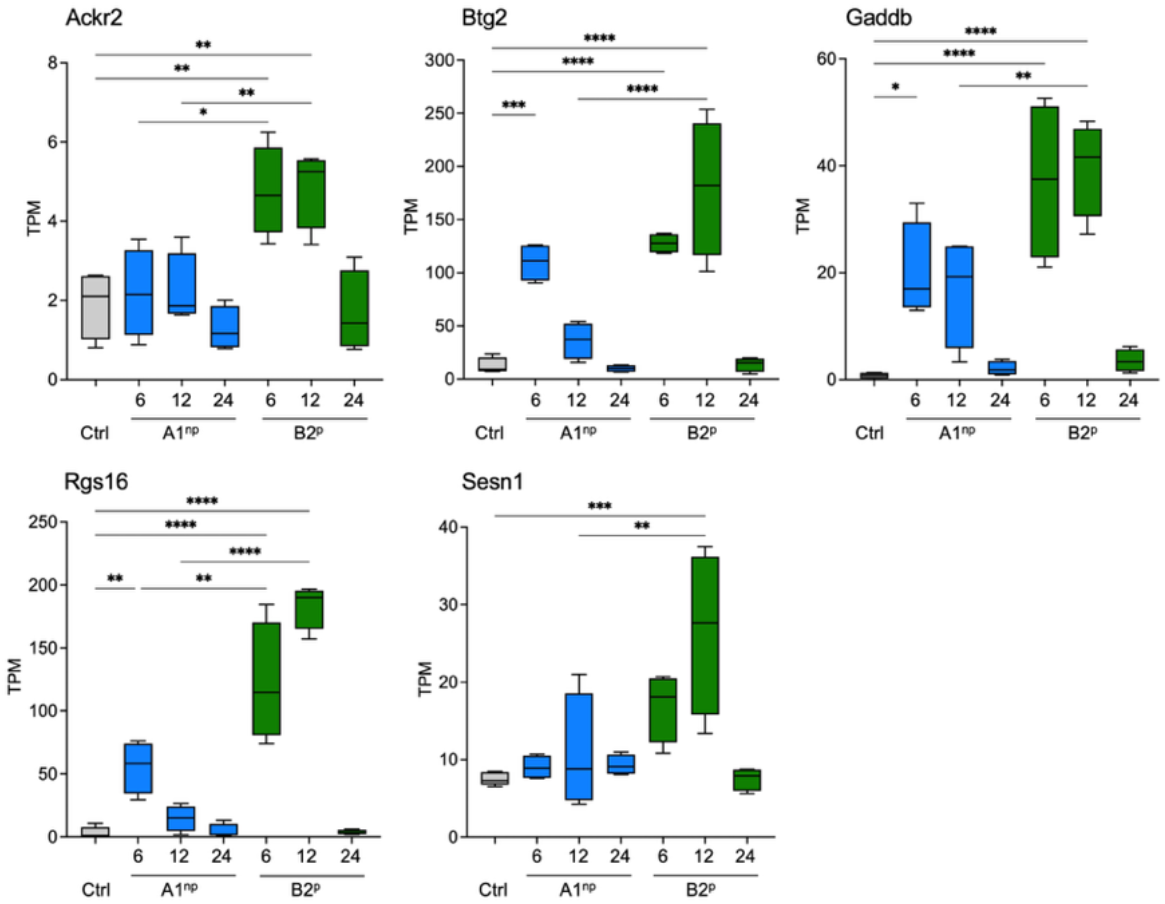
Time-dependent regulation of selected stress and immune response genes in the liver after infection with *Entamoeba histolytica*. Box plots show the transcript amounts (TPM) of *Ackr2, Btg2, Gadd45b, Rgs16* and *Sesn1* in liver tissue of untreated control animals (Ctrl, n = 5) and 6, 12 and 24 hours after intrahepatic infection with the non-pathogenic clone A1^np^ (blue, n = 4) or the pathogenic clone B2^p^ (green, n = 4). The data illustrate a pronounced, temporally differentiated gene induction, especially after B2^p^ infection, with strong activation of stress- and signal transduction-associated genes. Boxes represent median and interquartile range; whiskers represent minimum and maximum values. Statistical significance between groups is indicated by asterisks (* *p* < 0.05; ** *p* < 0.01; *** *p* < 0.001; **** p < 0.0001).

A direct comparison of these differentially expressed genes revealed distinct patterns of enriched biological processes at 12 and 24 hpi (Fig 7D-G).

At 12 hpi, genes that were upregulated in B2^p^-infected mice relative to A1^np^-infected mice were enriched in pathways related to cellular responses to oxygen-containing compounds, hormones, peptides and lipids (Fig 7D). Conversely, genes that were upregulated in A1^np^-relative to B2^p^-infected mice were associated with the regulation of metabolic processes, including primary and cellular metabolism, as well as DNA catabolic processes and cellular responses to DNA damage (Fig 7E).

At 24 hpi, genes that were upregulated in B2^p^-relative to A1^np^-infected mice were predominantly enriched in pathways related to the biosynthesis and metabolism of steroids, sterols, and cholesterol (Fig 7F). Conversely, genes that were upregulated in A1^np^-infected mice relative to B2^p^-infected mice were associated with triglyceride and acylglycerol metabolic processes, small molecule metabolism, and responses to hormonal stimuli (Fig 7G). Together, these data demonstrate time-dependent differences in the biological processes associated with differentially expressed genes in A1^np^ and B2^p^ infections.

### Validation of gene expression at the protein level reveals distinct expression and localization of immune regulators in amoebic liver abscesses

Since the liver damage is strongly mediated by B2^p^ trophozoites, infected liver tissue was stained for selected regulatory proteins that control inflammatory signals while preserving cellular integrity under stress. This analysis focused on a set of genes encoding proteins involved in chemokine regulation, inflammatory signalling, and cellular stress responses. Among them, Ackr2 (atypical chemokine receptor 2) acts as a chemokine-scavenging receptor that sequesters and degrades excessive chemokine levels [28, 29] thereby helping to prevent dysregulated chemokine-driven inflammation. Complementing this function, Rgs1 (regulator of G-protein signalling 1) serves as a negative regulator of pro-inflammatory immune responses [30], further contributing to the modulation of inflammatory signalling pathways. In addition to regulators of inflammation, several genes were identified that safeguard cellular homeostasis during stress. Btg2 (B-cell translocation gene 2) limits cell-cycle progression, supports controlled apoptosis, and contributes to DNA-damage responses, collectively preventing aberrant proliferation during normal tissue turnover [31]. Similarly, Gadd45b (Growth Arrest and DNA Damage–inducible 45 beta) is activated by environmental stresses, including genotoxic damage, to govern cell growth, survival, differentiation, and DNA repair [32]. Finally, Sesn1 (Sestrin 1) functions as a key sensor of cellular stress, particularly oxidative damage, and helps maintain redox balance [33].

To assess spatial distribution, localization analyses were exclusively performed on liver sections from mice infected with B2^p^ trophozoites. Immune cells that express *Ackr2, Btg2* and *Sesn1* appear to accumulate at the margin of the ALA. In the case of Gadd45b and Rgs16, the cells that present these molecules appear to migrate to the center of the abscess, where the trophozoites are also located (Fig 9).

**Fig. 9.**
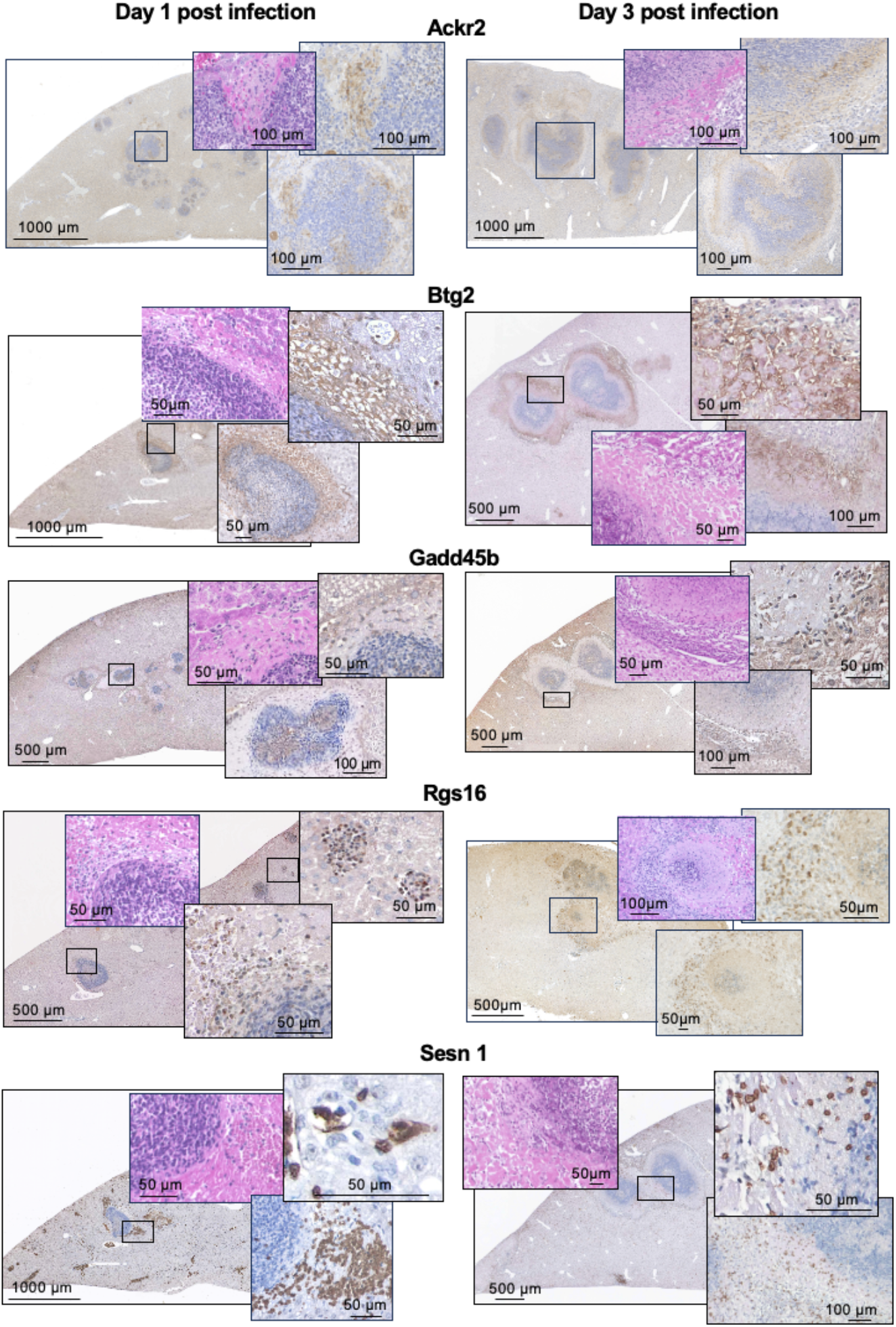
Localisation of Ackr2, Btg2, Gadd45b, Rgs16 and Sesn1 on immune cells one and three days after male mice were infected with B2^p^ trophozoites. Slices of paraffin-embedded liver sections from B2^p^ trophozoites-infected male mice obtained at day 1 and 3 post infection were stained with H&E or with polyclonal rabbit antibodies directed against Ackr2, Btg2, Gadd45b, Rgs16, and Sesn. Signal detection was carried out with an anti-rabbit HRP polymer system.

## Discussion

This study provides detailed insights into how two clones of *Entamoeba histolytica*, which differ in their pathogenicity, modulate the hepatic immune response. By comparing the transcriptional profiles of non-pathogenic A1^np^ and pathogenic B2^p^ trophozoites over time, we identified regulatory patterns that are both shared and distinct, and which shape the magnitude, composition and temporal dynamics of host immune activation.

A robust hepatic response was already evident at 6 hpi, with both trophozoite clones inducing substantial transcriptional changes. The number of differentially expressed genes peaked at 12 hpi and declined by 24 hpi, indicating transient but vigorous activation of liver immune signalling, which is consistent with previous reports [34, 35]. Notably, this difference was most pronounced at 12 hpi, where B2^p^ infection resulted in a markedly higher number of differentially expressed genes compared to A1^np^.

One limitation of this study is the absence of a sham-injected control group. As the intrahepatic injection procedure itself may induce local tissue injury and trigger early inflammatory and stress responses, it is possible that some of the observed transcriptional changes reflect effects related to the inoculation procedure rather than being parasite-specific. However, as both A1^np^- and B2^p^-infected mice underwent the same experimental treatment, it is possible to make a direct comparison between the two infection conditions. Therefore, although absolute transcriptional changes relative to uninfected controls may include some effects induced by the injection, the observed differences between A1^np^ and B2^p^ infections reflect clone-specific host responses that go beyond the shared procedural background.

Previous studies have primarily focused on infections involving B2^p^ trophozoites, demonstrating rapid infiltration of immune cells, particularly neutrophils and monocytes, into infected liver tissue. This contributes to both parasite control and tissue injury [2, 7]. Sellau *et al*. identified this cellular infiltration as a key driver of amoebic liver pathology [34]. Monocytes play a particularly important role, as an imbalance in their responses can disrupt immune homeostasis and promote pathological inflammation [2, 10]. In line with these observations, our data show that B2^p^ infection is associated with the sustained induction of chemokines such as *Ccl2, Ccl3, Ccl4, Cxcl1, Cxcl2*, and *Cxcl3*. This suggests the prolonged recruitment of innate immune cells. These findings are consistent with a more persistent inflammatory environment rather than a transient response.

Extracellular vesicles (EVs) derived from *E. histolytica* may contribute to liver pathology, as they have been shown to induce pro-inflammatory immune signalling in murine monocytes, including chemokine expression and activation of TNF pathways [6]. Notably, EVs from both A1^np^ and B2^p^ trophozoites have been reported to trigger such responses, suggesting that early immune activation may be a common feature of amoebic infection. In this context, the differences observed in our study are probably due to variations in the magnitude, persistence and downstream regulation of these responses, rather than to distinct initial activation mechanisms.

Transcriptomic analysis of the liver at 6 hpi, both A1^np^ and B2^p^ *E. histolytica* trophozoites revealed an increase of a conserved set of hepatic immune and stress responses. These responses are indicative of parasite recognition via pattern recognition receptors, such as Toll-like and cytokine receptors [36, 37]. This shared early response suggests activation of a common hepatic alarm programme. However, qualitative differences between the two infections were already apparent at this stage. Infection with A1^np^ trophozoites initiates a metabolically integrated immune programme characterized by the moderate activation of inflammatory and stress pathways within a regulatory context. This results in a stabilized cellular response that emphasizes metabolism, cytoprotection, and moderate immune activation. In contrast, B2^p^ trophozoites provoke a more differentiated and systemic immune response. This includes the activation of TLR receptor cascades alongside p53 and insulin signalling, increased interleukin activity, and early inflammatory amplification [38, 39]. This profile indicates an early shift toward enhanced inflammatory activation and cellular stress, consistent with the development of tissue pathology [34].

By 12 hpi, infection with the non-pathogenic A1^np^ clone was associated with the coordinated enrichment of pathways related to cytokine signalling, complement activation, and coagulation, as well as chemokine-mediated immune cell recruitment, as identified by KEGG and Reactome pathway analysis. These pathways are commonly linked to early innate immune responses and inflammatory regulation [40-42]. In addition, enrichment of IL-17 signalling pathways suggests the involvement of neutrophil-driven responses and acute inflammation, particularly at epithelial interfaces [43, 44]. Together with the observed modulation of lipid and steroid metabolism, these findings indicate a coordinated host response that integrates immune activation with metabolic adaptation, a pattern generally associated with the maintenance of tissue integrity during controlled inflammation [45, 46]. In contrast, infection with B2^p^ trophozoites, was characterised by a stronger activation of cell-intrinsic signalling pathways, including Toll-like receptor (TLR) pathways, notably TLR3 and TLR4, as well as robust cytokine signalling and activation of central downstream cascades, including NF-κB and MAPK [36, 47]. Enrichment of AGE-RAGE and TNF signalling pathways indicates oxidative stress, inflammation-driven apoptosis, and DNA damage [48, 49]. Furthermore, activation of mechanosensitive pathways, such as “fluid shear stress and atherosclerosis” suggests disturbances in vascular integrity and sinusoidal microarchitecture [50, 51]. These transcriptional changes are consistent with extensive metabolic reprogramming, cell cycle arrest, and the onset of insulin resistance, all of which are associated with tissue injury and disease progression [52]. The divergence between A1^np^ and B2^p^ infection appears to arise not from the presence or absence of these specific pathways, but rather from differences in their regulation, intensity, and integration within the overall host response. While A1^np^ drives a balanced and coordinated immune response, B2^p^ triggers a more pronounced and potentially dysregulated cellular stress response.

At 24 hpi, A1^np^ infection elicts to a balanced, non-pathogenic immune response associated with activation of metabolic and cytoprotective pathways, including those involved in lipid, steroid and vitamin A metabolism, as well as platelet-associated immunomodulatory processes [46, 53]. Overall, the response is controlled and non-escalatory, exhibiting regenerative features.

In comparison, B2^p^ infection at 24 hpi showed sustained activation of pro-inflammatory pathways, including cytokine and chemokine signalling, NF-κB, TNF, and complement cascades [54-57]. This persistent inflammatory state was accompanied by continued immune cell recruitment and a lack of compensatory detoxification responses, indicating ongoing tissue stress and damage [37, 58-60]. These findings are consistent with previous observations that liver regeneration occurs more rapidly following A1^np^ infection than after B2^p^ infection [18]. Sex-specific factors may further modulate these responses. Since hepatic amoebiasis predominantly affects males [61], all experiments in this study were performed using male mice [4]. HIF-1α has been identified as a key regulator of sex-dependent immune responses. It promotes Th17 polarisation while limiting Treg-mediated control, thereby enhancing inflammation-induced tissue damage [62]. In addition, testosterone has been shown to enhance the recruitment of neutrophils and monocytes via CXCL1, as well as modulating interferon responses and further amplifying inflammatory processes [7, 9, 34]. These mechanisms may contribute to the pronounced hyperinflammatory phenotype observed in B2^p^ infection.

To further dissect the molecular basis of these responses, we analysed the expression of immune-related genes. Although both A1^np^ and B2^p^ infection activated cytokine and chemokine signalling pathways, expression levels were generally higher following B2^p^ infection. Among the 151 immune-related genes, the majority displayed increased expression in B2^p^-infected livers, indicating stronger and more complex immune activation.

At 6 hpi, both groups exhibited upregulation of genes involved in pathogen recognition and inflammatory signalling, including *Clec4e* and *Cd14* as well as chemokines such as *Cxcl2*, the immunemetabolic regulator *Acod1* and the protease inhibitors *Serpina3n/m* [63-68].

Concurrent induction of regulatory genes such as *Cd300lf, Irak3*, and *Cd33* indicates early activation of feedback mechanisms limiting excessive inflammation [69-71].

However, distinct differences were already evident. A1^np^ infection showed stronger induction of acute-phase and antimicrobial genes, including *Serpine1, Lcn2*, and *Cxcl2*, suggesting rapid activation of innate defence mechanisms [72-75]. In contrast, B2^p^ infection resulted in heightened expression of genes promoting inflammation (*Osm, Rgs16, Igfbp1*), cellular stress responses (*Gadd45b, Gadd45g, Txnip*) and leukocyte recruitment (*Cxcl3, Mmp8*) [76-82].

At 12 hpi, both groups sustained activation of innate immune pathways, particularly those associated with neutrophil recruitment, as indicated by expression of *Cxcl1* and *Cxcr4* [83, 84]. Simultaneously, regulatory genes such as *Il1rn* and Serpina3m were upregulated, suggesting ongoing control of inflammatory damage [85, 86].

B2^p^ infection, however, was characterized by stronger induction of inflammatory and stress-related genes, including *Il17ra, Cxcl2, Ccr1*, and *Cxcr2*, and members of the *Gadd45* family [75, 81, 87-92]. This was accompanied by increased expression of *Il1r1, Tnfrsf1b, Phf11c*, and *Rgs16*, collectively indicating enhanced cytokine receptor signalling and immune cell activation [77, 93-96]. At the same time, regulatory genes associated with immune-dampening mechanisms, such as *Il1r2, Setdb2*, and *Ackr2*, were also upregulated, suggesting the concurrent activation of compensatory feedback pathways aimed at restraining excessive inflammation [97-100].

By 24 hpi, both groups exhibited signs of immune resolution, stabilised by anti-inflammatory signalling and tissue repair. This included the induction of anti-inflammatory and regulatory genes, such as *Hp, Cish, Cd33* and *Gas6*, as well as the coordinated regulation of genes involved in immune cell trafficking, maturation and adhesion (*Cxcr4, Itgam, Csf3r*) [84, 101-107].

However, B2^p^ infection was characterised by markedly elevated expression of pro-inflammatory and acute-phase mediators, including *Cxcl2, Cxcl3, Ccl3, Saa3, Osm, Ptgs2* and *Trem1*. This indicates sustained immune activation and ongoing tissue stress [73, 75, 76, 79, 108, 109]. The persistently high expression of *Il1r2* suggested sustained anti-inflammatory feedback. Meanwhile, the elevated levels of *Serpine1* possibly contributed to tissue protection and the resolution of inflammation [97, 98].

In our study, immunohistochemical staining revealed distinct spatial patterns of cells expressing genes that were strongly up-regulated during B2^p^ infection. Immune cells positive for Ackr2, Btg2 and Sesn1 accumulated predominantly at the periphery of the amoebic abscess. In contrast, cells expressing Gadd45b and Rgs16 localised to the abscess core, coming into close proximity with the trophozoites. This distribution likely reflects the distinct functional programmes of the infiltrating immune cells. Cells at the abscess margin appear to engage primarily in regulatory and protective activities. Ackr2 limits chemokine-driven recruitment, Btg2 restrains lymphocyte overactivation and Sesn1 reduces reactive oxygen species and metabolic stress [31, 100, 110-113]. Together, these factors may stabilise the inflammatory environment, preventing excessive tissue damage while containing the lesion. In contrast, immune cells expressing Gadd45b and Rgs16 were enriched at the parasite–host interface. Gadd45b is associated with stress-induced signalling and effector activation, while Rgs16 modulates GPCR-driven chemokine responses, facilitating the precise positioning of effector cells [77, 89]. Their central localisation suggests a role in direct antiparasitic defence and tissue stress adaptation, which is consistent with the requirement for strong effector functions at sites where trophozoites are present.

Overall, these findings support the idea that immune programmes are spatially segregated during amoebic liver abscess formation, with regulatory responses at the periphery and stress-adapted effector functions at the centre. This represents a strategy that balances parasite control with limiting collateral tissue damage.

## Conclusion

This study reveals that the immune response to *E. histolytica* in the liver is shaped by both conserved core pathways and clone-specific dynamics. While both clones activate innate defences, including cytokines, chemokines and regulatory feedback, the non-pathogenic A1^np^ clone triggers a rapid but self-limiting inflammatory response involving antimicrobial defence and tissue protection. By contrast, the pathogenic B2^p^ clone provokes a more sustained and multifaceted pro-inflammatory immune response characterised by oxidative stress, cytokine amplification and tissue remodelling. These findings suggest that immune dysregulation in response to distinct parasitic clones, rather than parasite presence alone, is a key driver of liver pathology in ALA. Understanding these divergent immune profiles could inform future therapies aimed at mitigating tissue damage by modulating host responses without compromising pathogen control.

## Acknowledgment

We would like to thank Susann Ofori for her technical assistance and excellent laboratory management.

## Author contribution

**Conceptualization:** Helena Fehling, Hanna Lotter, Iris Bruchhaus.

**Data curation:** Johannes Allweier, Helena Fehling, Iris Bruchhaus

**Formal analysis:** Helena Fehling, Johannes Allweier, Barbara Honecker, Iris Bruchhaus.

**Funding acquisition:** Hanna Lotter, Iris Bruchhaus.

**Investigation:** Helena Fehling, Johannes Allweier, Claudia Marggraff.

**Methodology:** Helena Fehling, Claudia Marggraff.

**Supervision:** Hanna Lotter, Iris Bruchhaus.

**Visualization:** Barbara Honecker, Johannes Allweier, Iris Bruchhaus.

**Writing – original draft:** Iris Bruchhaus.

**Writing – review & editing:** Helena Fehling, Johannes Allweier, Barbara Honecker, Juliett Anders, Michel-Ruben Glagowski, Hanna Lotter, Iris Bruchhaus

## Data Availability Statement

All relevant data are within the manuscript and its Supporting information files. The raw data were submitted to NCBI-GEO (Gene Expression Omnibus) under the series record GSE329300 and are available as expression browser in Table S1.

## References

1. Collaborators GBDDD. Estimates of the global, regional, and national morbidity, mortality, and aetiologies of diarrhoea in 195 countries: a systematic analysis for the Global Burden of Disease Study 2016. Lancet Infect Dis. 2018;18(11):1211–28. Epub 2018/09/24. doi: 10.1016/S1473-3099(18)30362-1. PMID: 30243583.

2. Helk E, Bernin H, Ernst T, Ittrich H, Jacobs T, Heeren J, et al. TNFalpha-mediated liver destruction by Kupffer cells and Ly6Chi monocytes during Entamoeba histolytica infection. PLoS pathog. 2013;9(1):e1003096. Epub 2013/01/10. doi: 10.1371/journal.ppat.1003096. PMID: 23300453.

3. Estrada-Figueroa LA, Ramirez-Jimenez Y, Osorio-Trujillo C, Shibayama M, Navarro-Garcia F, Garcia-Tovar C, et al. Absence of CD38 delays arrival of neutrophils to the liver and innate immune response development during hepatic amoebiasis by Entamoeba histolytica. Parasite Immunol. 2011;33(12):661–8. doi: 10.1111/j.1365-3024.2011.01333.x. PMID: 21919917.

4. Lotter H, Jacobs T, Gaworski I, Tannich E. Sexual dimorphism in the control of amebic liver abscess in a mouse model of disease. Infection and immunity. 2006;74(1):118–24. PMID: 16368964.

5. Pacheco-Yepez J, Galvan-Moroyoqui JM, Meza I, Tsutsumi V, Shibayama M. Expression of cytokines and their regulation during amoebic liver abscess development. Parasite Immunol. 2011;33(1):56–64. doi: 10.1111/j.1365-3024.2010.01252.x PMID: 21155843.

6. Honecker B, Barreiter VA, Hohn K, Horvath B, Harant K, Metwally NG, et al. Entamoeba histolytica extracellular vesicles drive pro-inflammatory monocyte signaling. PLoS Negl Trop Dis. 2025;19(4):e0012997. Epub 20250410. doi: 10.1371/journal.pntd.0012997 PMID: 40208874.

7. Er-Lukowiak M, Hanzelmann S, Rothe M, Moamenpour DT, Hausmann F, Khatri R, et al. Testosterone affects type I/type II interferon response of neutrophils during hepatic amebiasis. Front Immunol. 2023;14:1279245. Epub 20231221. doi: 10.3389/fimmu.2023.1279245. PMID: 38179044.

8. Garcia-Zepeda EA, Rojas-Lopez A, Esquivel-Velazquez M, Ostoa-Saloma P. Regulation of the inflammatory immune response by the cytokine/chemokine network in amoebiasis. Parasite Immunol. 2007;29(12):679–84. doi: 10.1111/j.1365-3024.2007.00990.x. PMID: 18042174.

9. Sellau J, Groneberg M, Fehling H, Thye T, Hoenow S, Marggraff C, et al. Androgens predispose males to monocyte-mediated immunopathology by inducing the expression of leukocyte recruitment factor CXCL1. Nat Commun. 2020;11(1):3459. Epub 20200710. doi: 10.1038/s41467-020-17260-y. PMID: 32651360.

10. Sellau J, Puengel T, Hoenow S, Groneberg M, Tacke F, Lotter H. Monocyte dysregulation: consequences for hepatic infections. Semin Immunopathol. 2021;43(4):493–506. Epub 20210407. doi: 10.1007/s00281-021-00852-1. PMID: 33829283.

11. Maldonado-Bernal C, Kirschning CJ, Rosenstein Y, Rocha LM, Rios-Sarabia N, Espinosa-Cantellano M, et al. The innate immune response to Entamoeba histolytica lipopeptidophosphoglycan is mediated by toll-like receptors 2 and 4. Parasite Immunol. 2005;27(4):127–37. doi: 10.1111/j.1365-3024.2005.00754.x. PMID: 15910421.

12. Faust DM, Marquay Markiewicz J, Santi-Rocca J, Guillen N. New insights into host-pathogen interactions during Entamoeba histolytica liver infection. Eur J Microbiol Immunol (Bp). 2011;1(1):10–8. doi: 10.1556/EuJMI.1.2011.1.4. PMID: 24466432.

13. Begum S, Quach J, Chadee K. Immune Evasion Mechanisms of Entamoeba histolytica: Progression to Disease. Front Microbiol. 2015;6:1394. Epub 20151215. doi: 10.3389/fmicb.2015.01394. PMID: 26696997.

14. Fonseca Z, Diaz-Godinez C, Mora N, Aleman OR, Uribe-Querol E, Carrero JC, et al. Entamoeba histolytica Induce Signaling via Raf/MEK/ERK for Neutrophil Extracellular Trap (NET) Formation. Front Cell Infect Microbiol. 2018;8:226. Epub 20180704. doi: 10.3389/fcimb.2018.00226. PMID: 30023352.

15. Avila EE, Salaiza N, Pulido J, Rodriguez MC, Diaz-Godinez C, Laclette JP, et al. Entamoeba histolytica Trophozoites and Lipopeptidophosphoglycan Trigger Human Neutrophil Extracellular Traps. PLoS One. 2016;11(7):e0158979. Epub 20160714. doi: 10.1371/journal.pone.0158979. PMID: 27415627.

16. Roy M, Chakraborty S, Kumar Srivastava S, Kaushik S, Jyoti A, Kumar Srivastava V. Entamoeba histolytica induced NETosis and the dual role of NETs in amoebiasis. Int Immunopharmacol. 2023;118:110100. Epub 20230401. doi: 10.1016/j.intimp.2023.110100. PMID: 37011501.

17. Diaz-Godinez C, Rios-Valencia DG, Garcia-Aguirre S, Martinez-Calvillo S, Carrero JC. Immunomodulatory effect of extracellular vesicles from Entamoeba histolytica trophozoites: Regulation of NETs and respiratory burst during confrontation with human neutrophils. Front Cell Infect Microbiol. 2022;12:1018314. Epub 20221028. doi: 10.3389/fcimb.2022.1018314. PMID: 36389143.

18. Meyer M, Fehling H, Matthiesen J, Lorenzen S, Schuldt K, Bernin H, et al. Overexpression of differentially expressed genes identified in non-pathogenic and pathogenic Entamoeba histolytica clones allow identification of new pathogenicity factors involved in amoebic liver abscess formation. PLoS Pathog. 2016;12(8):e1005853. doi: 10.1371/journal.ppat.1005853. PMID: 27575775.

19. Konig C, Honecker B, Wilson IW, Weedall GD, Hall N, Roeder T, et al. Taxon-Specific Proteins of the Pathogenic Entamoeba Species E. histolytica and E. nuttalli. Front Cell Infect Microbiol. 2021;11:641472. Epub 20210319. doi: 10.3389/fcimb.2021.641472. PMID: 33816346.

20. Petri WA, Jr., Schnaar RL. Purification and characterization of galactose- and N-acetylgalactosamine-specific adhesin lectin of Entamoeba histolytica. Methods Enzymol. 1995;253:98–104.

21. Matthiesen J, Bar AK, Bartels AK, Marien D, Ofori S, Biller L, et al. Overexpression of specific cysteine peptidases confers pathogenicity to a nonpathogenic Entamoeba histolytica clone. MBio. 2013;4(2):e00072–13. Epub 2013/03/28. doi: 10.1128/mBio.00072-13. PMID: 23532975.

22. Leippe M. Amoebapores. Parasitol Today. 1997;13(5):178–83.

23. Matthiesen J, Lender C, Haferkorn A, Fehling H, Meyer M, Matthies T, et al. Trigger-induced RNAi gene silencing to identify pathogenicity factors of Entamoeba histolytica. FASEB J. 2019;33(2):1658–68. doi: 10.1096/fj.201801313R. PMID: 30169111.

24. Diamond LS, Harlow DR, Cunnick CC. A new medium for the axenic cultivation of Entamoeba histolytica and other Entamoeba. Trans R Soc Trop Med Hyg. 1978;72(4):431–2. doi: 10.1016/0035-9203(78)90144-x. PMID: 212851.

25. Babicki S, Arndt D, Marcu A, Liang Y, Grant JR, Maciejewski A, et al. Heatmapper: web-enabled heat mapping for all. Nucleic Acids Res. 2016;44(W1):W147–53. Epub 20160517. doi: 10.1093/nar/gkw419. PMID: 27190236.

26. Szklarczyk D, Kirsch R, Koutrouli M, Nastou K, Mehryary F, Hachilif R, et al. The STRING database in 2023: protein-protein association networks and functional enrichment analyses for any sequenced genome of interest. Nucleic Acids Res. 2023;51(D1):D638–D46. doi: 10.1093/nar/gkac1000. PMID: 36370105.

27. Galaxy C. The Galaxy platform for accessible, reproducible, and collaborative data analyses: 2024 update. Nucleic Acids Res. 2024;52(W1):W83–W94. doi: 10.1093/nar/gkae410. PMID: 38769056.

28. Massara M, Bonavita O, Savino B, Caronni N, Mollica Poeta V, Sironi M, et al. ACKR2 in hematopoietic precursors as a checkpoint of neutrophil release and anti-metastatic activity. Nat Commun. 2018;9(1):676. Epub 20180214. doi: 10.1038/s41467-018-03080-8. PMID: 29445158.

29. Bonavita O, Mollica Poeta V, Setten E, Massara M, Bonecchi R. ACKR2: An Atypical Chemokine Receptor Regulating Lymphatic Biology. Front Immunol. 2016;7:691. Epub 20170111. doi: 10.3389/fimmu.2016.00691. PMID: 28123388.

30. Suurvali J, Pahtma M, Saar R, Paalme V, Nutt A, Tiivel T, et al. RGS16 restricts the pro-inflammatory response of monocytes. Scand J Immunol. 2015;81(1):23–30. doi: 10.1111/sji.12250. PMID: 25366993.

31. Kim SH, Jung IR, Hwang SS. Emerging role of anti-proliferative protein BTG1 and BTG2. BMB Rep. 2022;55(8):380–8. doi: 10.5483/BMBRep.2022.55.8.092. PMID: 35880434.

32. Rodriguez-Jimenez P, Fernandez-Messina L, Ovejero-Benito MC, Chicharro P, Vera-Tome P, Vara A, et al. Growth arrest and DNA damage-inducible proteins (GADD45) in psoriasis. Sci Rep. 2021;11(1):14579. Epub 20210716. doi: 10.1038/s41598-021-93780-x. PMID: 34272424.

33. Ro SH, Fay J, Cyuzuzo CI, Jang Y, Lee N, Song HS, et al. SESTRINs: Emerging Dynamic Stress-Sensors in Metabolic and Environmental Health. Front Cell Dev Biol. 2020;8:603421. Epub 20201203. doi: 10.3389/fcell.2020.603421. PMID: 33425907.

34. Sellau J, Groneberg M, Hoenow S, Lotter H. The underlying cellular immune pathology of Entamoeba histolytica-induced hepatic amoebiasis. J Hepatol. 2021;75(2):481–2. Epub 20210611. doi: 10.1016/j.jhep.2021.03.018. PMID: 34120776.

35. Pulido-Ortega J, Talamas-Rohana P, Munoz-Ortega MH, Aldaba-Muruato LR, Martinez-Hernandez SL, Campos-Esparza MDR, et al. Functional Characterization of an Interferon Gamma Receptor-Like Protein on Entamoeba histolytica. Infection and immunity. 2019;87(11). Epub 20191018. doi: 10.1128/IAI.00540-19. PMID: 31427448.

36. Duan T, Du Y, Xing C, Wang HY, Wang RF. Toll-Like Receptor Signaling and Its Role in Cell-Mediated Immunity. Front Immunol. 2022;13:812774. Epub 20220303. doi: 10.3389/fimmu.2022.812774. PMID: 35309296.

37. Kiziltas S. Toll-like receptors in pathophysiology of liver diseases. World J Hepatol. 2016;8(32):1354–69. doi: 10.4254/wjh.v8.i32.1354. PMID: 27917262.

38. Chen L, Deng H, Cui H, Fang J, Zuo Z, Deng J, et al. Inflammatory responses and inflammation-associated diseases in organs. Oncotarget. 2018;9(6):7204–18. Epub 20171214. doi: 10.18632/oncotarget.23208. PMID: 29467962.

39. Salauddin M, Bhattacharyya D, Samanta I, Saha S, Xue M, Hossain MG, et al. Role of TLRs as signaling cascades to combat infectious diseases: a review. Cell Mol Life Sci. 2025;82(1):122. Epub 20250319. doi: 10.1007/s00018-025-05631-x. PMID: 40105962.

40. Huber-Lang M, Kovtun A, Ignatius A. The role of complement in trauma and fracture healing. Semin Immunol. 2013;25(1):73–8. Epub 20130613. doi: 10.1016/j.smim.2013.05.006. PMID: 23768898.

41. Czermak BJ, Lentsch AB, Bless NM, Schmal H, Friedl HP, Ward PA. Synergistic enhancement of chemokine generation and lung injury by C5a or the membrane attack complex of complement. Am J Pathol. 1999;154(5):1513–24. doi: 10.1016/S0002-9440(10)65405-3. PMID: 10329604.

42. Diaz-Valencia JD, Perez-Yepez EA, Ayala-Sumuano JT, Franco E, Meza I. A surface membrane protein of Entamoeba histolytica functions as a receptor for human chemokine IL-8: its role in the attraction of trophozoites to inflammation sites. Int J Parasitol. 2015;45(14):915–23. Epub 20150904. doi: 10.1016/j.ijpara.2015.07.007. PMID: 26343219.

43. Paquissi FC. Immune Imbalances in Non-Alcoholic Fatty Liver Disease: From General Biomarkers and Neutrophils to Interleukin-17 Axis Activation and New Therapeutic Targets. Front Immunol. 2016;7:490. Epub 20161111. doi: 10.3389/fimmu.2016.00490. PMID: 27891128.

44. Shen H, Liangpunsakul S, Iwakiri Y, Szabo G, Wang H. Immunological mechanisms and emerging therapeutic targets in alcohol-associated liver disease. Cell Mol Immunol. 2025. Epub 20250521. doi: 10.1038/s41423-025-01291-w. PMID: 40399593.

45. Rudnick DA, Davidson NO. Functional Relationships between Lipid Metabolism and Liver Regeneration. Int J Hepatol. 2012;2012:549241. Epub 20120126. doi: 10.1155/2012/549241. PMID: 22319652.

46. Garcia C, Andersen CJ, Blesso CN. The Role of Lipids in the Regulation of Immune Responses. Nutrients. 2023;15(18). Epub 20230907. doi: 10.3390/nu15183899. PMID: 37764683.

47. Kawai T, Ikegawa M, Ori D, Akira S. Decoding Toll-like receptors: Recent insights and perspectives in innate immunity. Immunity. 2024;57(4):649–73. doi: 10.1016/j.immuni.2024.03.004. PMID: 38599164.

48. Zhou M, Zhang Y, Shi L, Li L, Zhang D, Gong Z, et al. Activation and modulation of the AGEs-RAGE axis: Implications for inflammatory pathologies and therapeutic interventions - A review. Pharmacol Res. 2024;206:107282. Epub 20240622. doi: 10.1016/j.phrs.2024.107282. PMID: 38914383.

49. Yue Q, Song Y, Liu Z, Zhang L, Yang L, Li J. Receptor for Advanced Glycation End Products (RAGE): A Pivotal Hub in Immune Diseases. Molecules. 2022;27(15). Epub 20220802. doi: 10.3390/molecules27154922. PMID: 35956875.

50. Cheng H, Zhong W, Wang L, Zhang Q, Ma X, Wang Y, et al. Effects of shear stress on vascular endothelial functions in atherosclerosis and potential therapeutic approaches. Biomed Pharmacother. 2023;158:114198. Epub 20230103. doi: 10.1016/j.biopha.2022.114198. PMID: 36916427.

51. Conway DE, Schwartz MA. Flow-dependent cellular mechanotransduction in atherosclerosis. J Cell Sci. 2013;126(Pt 22):5101–9. Epub 20131104. doi: 10.1242/jcs.138313. PMID: 24190880.

52. Malhi H, Gores GJ. Cellular and molecular mechanisms of liver injury. Gastroenterology. 2008;134(6):1641–54. doi: 10.1053/j.gastro.2008.03.002. PMID: 18471544.

53. Lisman T, Luyendyk JP. Platelets as Modulators of Liver Diseases. Semin Thromb Hemost. 2018;44(2):114–25. Epub 20170912. doi: 10.1055/s-0037-1604091. PMID: 28898899.

54. Tak PP, Firestein GS. NF-kappaB: a key role in inflammatory diseases. J Clin Invest. 2001;107(1):7–11. doi: 10.1172/JCI11830. PMID: 11134171.

55. Bhol NK, Bhanjadeo MM, Singh AK, Dash UC, Ojha RR, Majhi S, et al. The interplay between cytokines, inflammation, and antioxidants: mechanistic insights and therapeutic potentials of various antioxidants and anti-cytokine compounds. Biomed Pharmacother. 2024;178:117177. Epub 20240724. doi: 10.1016/j.biopha.2024.117177. PMID: 39053423.

56. Dorrington MG, Fraser IDC. NF-kappaB Signaling in Macrophages: Dynamics, Crosstalk, and Signal Integration. Front Immunol. 2019;10:705. Epub 20190409. doi: 10.3389/fimmu.2019.00705. PMID: 31024544.

57. Tacke F, Luedde T, Trautwein C. Inflammatory pathways in liver homeostasis and liver injury. Clin Rev Allergy Immunol. 2009;36(1):4–12. doi: 10.1007/s12016-008-8091-0. PMID: 18600481.

58. Robinson MW, Harmon C, O’Farrelly C. Liver immunology and its role in inflammation and homeostasis. Cell Mol Immunol. 2016;13(3):267–76. Epub 20160411. doi: 10.1038/cmi.2016.3. PMID: 27063467.

59. Martin-Mateos R, Alvarez-Mon M, Albillos A. Dysfunctional Immune Response in Acute-on-Chronic Liver Failure: It Takes Two to Tango. Front Immunol. 2019;10:973. Epub 20190501. doi: 10.3389/fimmu.2019.00973. PMID: 31118937.

60. Nakamoto N, Kanai T. Role of toll-like receptors in immune activation and tolerance in the liver. Front Immunol. 2014;5:221. Epub 20140516. doi: 10.3389/fimmu.2014.00221. PMID: 24904576.

61. Blessmann J, Van Linh P, Nu PA, Thi HD, Muller-Myhsok B, Buss H, et al. Epidemiology of amebiasis in a region of high incidence of amebic liver abscess in central Vietnam. Am J Trop Med Hyg. 2002;66(5):578–83. doi: 10.4269/ajtmh.2002.66.578. PMID: 12201594.

62. Groneberg M, Hoenow S, Marggraff C, Fehling H, Metwally NG, Hansen C, et al. HIF-1alpha modulates sex-specific Th17/Treg responses during hepatic amoebiasis. J Hepatol. 2022;76(1):160–73. Epub 20210929. doi: 10.1016/j.jhep.2021.09.020. PMID: 34599999.

63. Pahari S, Negi S, Aqdas M, Arnett E, Schlesinger LS, Agrewala JN. Induction of autophagy through CLEC4E in combination with TLR4: an innovative strategy to restrict the survival of Mycobacterium tuberculosis. Autophagy. 2020;16(6):1021–43. Epub 20190908. doi: 10.1080/15548627.2019.1658436. PMID: 31462144.

64. Na K, Oh BC, Jung Y. Multifaceted role of CD14 in innate immunity and tissue homeostasis. Cytokine Growth Factor Rev. 2023;74:100–7. Epub 20230823. doi: 10.1016/j.cytogfr.2023.08.008. PMID: 37661484.

65. Ding L, Hayes MM, Photenhauer A, Eaton KA, Li Q, Ocadiz-Ruiz R, et al. Schlafen 4-expressing myeloid-derived suppressor cells are induced during murine gastric metaplasia. J Clin Invest. 2016;126(8):2867–80. Epub 20160718. doi: 10.1172/JCI82529. PMID: 27427984.

66. De Filippo K, Henderson RB, Laschinger M, Hogg N. Neutrophil chemokines KC and macrophage-inflammatory protein-2 are newly synthesized by tissue macrophages using distinct TLR signaling pathways. J Immunol. 2008;180(6):4308–15. doi: 10.4049/jimmunol.180.6.4308. PMID: 18322244.

67. Wu R, Liu J, Tang D, Kang R. The Dual Role of ACOD1 in Inflammation. J Immunol. 2023;211(4):518–26. doi: 10.4049/jimmunol.2300101. PMID: 37549395.

68. de Mezer M, Rogalinski J, Przewozny S, Chojnicki M, Niepolski L, Sobieska M, et al. SERPINA3: Stimulator or Inhibitor of Pathological Changes. Biomedicines. 2023;11(1). Epub 20230107. doi: 10.3390/biomedicines11010156. PMID: 36672665.

69. Sutherland SIM, Ju X, Silveira PA, Kupresanin F, Horvath LG, Clark GJ. CD300f signalling induces inhibitory human monocytes/macrophages. Cell Immunol. 2023;390:104731. Epub 20230602. doi: 10.1016/j.cellimm.2023.104731. PMID: 37302321.

70. Freihat LA, Wheeler JI, Wong A, Turek I, Manallack DT, Irving HR. IRAK3 modulates downstream innate immune signalling through its guanylate cyclase activity. Sci Rep. 2019;9(1):15468. Epub 20191029. doi: 10.1038/s41598-019-51913-3. PMID: 31664109.

71. Lajaunias F, Dayer JM, Chizzolini C. Constitutive repressor activity of CD33 on human monocytes requires sialic acid recognition and phosphoinositide 3-kinase-mediated intracellular signaling. Eur J Immunol. 2005;35(1):243–51. doi: 10.1002/eji.200425273. PMID: 15597323.

72. Kubota M, Yoshida Y, Kobayashi E, Matsutani T, Li SY, Zhang BS, et al. Serum anti-SERPINE1 antibody as a potential biomarker of acute cerebral infarction. Sci Rep. 2021;11(1):21772. Epub 20211105. doi: 10.1038/s41598-021-01176-8. MID: 34741085.

73. Brusletto BS, Loberg EM, Hellerud BC, Goverud IL, Berg JP, Olstad OK, et al. Extensive Changes in Transcriptomic “Fingerprints” and Immunological Cells in the Large Organs of Patients Dying of Acute Septic Shock and Multiple Organ Failure Caused by Neisseria meningitidis. Front Cell Infect Microbiol. 2020;10:42. Epub 20200219. doi: 10.3389/fcimb.2020.00042. PMID: 32154187.

74. Chang YL, Wang Z, Igawa S, Choi JE, Werbel T, Di Nardo A. Lipocalin 2: A New Antimicrobial in Mast Cells. Int J Mol Sci. 2019;20(10). Epub 20190514. doi: 10.3390/ijms20102380. PMID: 31091692.

75. Lv Y, Chen C, Han M, Tian C, Song F, Feng S, et al. CXCL2: a key player in the tumor microenvironment and inflammatory diseases. Cancer Cell Int. 2025;25(1):133. Epub 20250407. doi: 10.1186/s12935-025-03765-3. PMID: 40197328.

76. Wolf CL, Pruett C, Lighter D, Jorcyk CL. The clinical relevance of OSM in inflammatory diseases: a comprehensive review. Front Immunol. 2023;14:1239732. Epub 20230929. doi: 10.3389/fimmu.2023.1239732. PubMed PMID: 37841259; PubMed Central PMCID: PMCPMC10570509.

77. Zhu F, Qin Y, Wang Y, Zhang F, Xu Z, Dai F, et al. Critical roles of RGS16 in the mucosal inflammation of ulcerative colitis. Eur J Gastroenterol Hepatol. 2022;34(10):993–9. Epub 20220805. doi: 10.1097/MEG.0000000000002407. PMID: 35830366.

78. Lewitt MS, Boyd GW. Insulin-like Growth Factor-Binding Protein-1 (IGFBP-1) as a Biomarker of Cardiovascular Disease. Biomolecules. 2024;14(11). Epub 20241120. doi: 10.3390/biom14111475. PMID: 39595651.

79. Bao Y, Tong C, Xiong X. CXCL3: A key player in tumor microenvironment and inflammatory diseases. Life Sci. 2024;348:122691. Epub 20240505. doi: 10.1016/j.lfs.2024.122691. PubMed PMID: 38714265.

80. Khatwa UA, Kleibrink BE, Shapiro SD, Subramaniam M. MMP-8 promotes polymorphonuclear cell migration through collagen barriers in obliterative bronchiolitis. J Leukoc Biol. 2010;87(1):69–77. Epub 20091002. doi: 10.1189/jlb.0509361. PMID: 19801498.

81. Liebermann DA, Hoffman B. Gadd45 in stress signaling. J Mol Signal. 2008;3:15. Epub 20080912. doi: 10.1186/1750-2187-3-15. PMID: 18789159.

82. Choi EH, Park SJ. TXNIP: A key protein in the cellular stress response pathway and a potential therapeutic target. Exp Mol Med. 2023;55(7):1348–56. Epub 20230703. doi: 10.1038/s12276-023-01019-8. PMID: 37394581.

83. Sawant KV, Poluri KM, Dutta AK, Sepuru KM, Troshkina A, Garofalo RP, et al. Chemokine CXCL1 mediated neutrophil recruitment: Role of glycosaminoglycan interactions. Sci Rep. 2016;6:33123. Epub 20160914. doi: 10.1038/srep33123. PMID: 27625115.

84. De Filippo K, Rankin SM. CXCR4, the master regulator of neutrophil trafficking in homeostasis and disease. Eur J Clin Invest. 2018;48 Suppl 2(Suppl Suppl 2):e12949. Epub 20180523. doi: 10.1111/eci.12949. PMID: 29734477.

85. Zhang M, Huang Y, Pan J, Sang C, Lin Y, Dong L, et al. An Inflammatory Checkpoint Generated by IL1RN Splicing Offers Therapeutic Opportunity for KRAS-Mutant Intrahepatic Cholangiocarcinoma. Cancer Discov. 2023;13(10):2248–69. doi: 10.1158/2159-8290.CD-23-0282. PMID: 37486241.

86. Heit C, Jackson BC, McAndrews M, Wright MW, Thompson DC, Silverman GA, et al. Update of the human and mouse SERPIN gene superfamily. Hum Genomics. 2013;7(1):22. Epub 20131030. doi: 10.1186/1479-7364-7-22. PMID: 24172014.

87. Ge S, Hertel B, Susnik N, Rong S, Dittrich AM, Schmitt R, et al. Interleukin 17 receptor A modulates monocyte subsets and macrophage generation in vivo. PLoS One. 2014;9(1):e85461. Epub 20140115. doi: 10.1371/journal.pone.0085461. PMID: 24454873.

88. Zimmermann HW, Tacke F. Modification of chemokine pathways and immune cell infiltration as a novel therapeutic approach in liver inflammation and fibrosis. Inflamm Allergy Drug Targets. 2011;10(6):509–36. doi: 10.2174/187152811798104890. PMID: 22150762.

89. Salerno DM, Tront JS, Hoffman B, Liebermann DA. Gadd45a and Gadd45b modulate innate immune functions of granulocytes and macrophages by differential regulation of p38 and JNK signaling. J Cell Physiol. 2012;227(11):3613–20. doi: 10.1002/jcp.24067. PMID: 22307729.

90. Takekawa M, Saito H. A family of stress-inducible GADD45-like proteins mediate activation of the stress-responsive MTK1/MEKK4 MAPKKK. Cell. 1998;95(4):521–30. doi: 10.1016/s0092-8674(00)81619-0. PMID: 9827804.

91. Dostie KE, Thees AV, Lynes MA. Metallothionein: A Novel Therapeutic Target for Treatment of Inflammatory Bowel Disease. Curr Pharm Des. 2018;24(27):3155–61. doi: 10.2174/1381612824666180717110236. PMID: 30014800.

92. Devisscher L, Hindryckx P, Lynes MA, Waeytens A, Cuvelier C, De Vos F, et al. Role of metallothioneins s danger signals in the pathogenesis of colitis. J Pathol. 2014;233(1):89–100. Epub 20140224. doi: 10.1002/path.4330. PMID: 24452846.

93. Auron PE. The interleukin 1 receptor: ligand interactions and signal transduction. Cytokine Growth Factor Rev. 1998;9(3-4):221–37. doi: 10.1016/s1359-6101(98)00018-5. PMID: 9918122.

94. Rahman N, Stewart G, Jones G. A role for the atopy-associated gene PHF11 in T-cell activation and viability. Immunol Cell Biol. 2010;88(8):817–24. Epub 20100427. doi: 10.1038/icb.2010.57. PMID: 20421878.

95. Gao Y, Shi H, Zhao H, Yao M, He Y, Jiang M, et al. Single-cell transcriptomics identify TNFRSF1B as a novel T-cell exhaustion marker for ovarian cancer. Clin Transl Med. 2023;13(9):e1416. doi: 10.1002/ctm2.1416. PMID: 37712139.

96. McCulloch TR, Rossi GR, Alim L, Lam PY, Wong JKM, Coleborn E, et al. Dichotomous outcomes of TNFR1 and TNFR2 signaling in NK cell-mediated immune responses during inflammation. Nat Commun. 2024;15(1):9871. Epub 20241114. doi: 10.1038/s41467-024-54232-y. PMID: 39543125.

97. Supino D, Minute L, Mariancini A, Riva F, Magrini E, Garlanda C. Negative Regulation of the IL-1 System by IL-1R2 and IL-1R8: Relevance in Pathophysiology and Disease. Front Immunol. 2022;13:804641. Epub 20220208. doi: 10.3389/fimmu.2022.804641. PMID: 35211118.

98. Molgora M, Supino D, Mantovani A, Garlanda C. Tuning inflammation and immunity by the negative regulators IL-1R2 and IL-1R8. Immunol Rev. 2018;281(1):233–47. doi: 10.1111/imr.12609. PMID: 29247989.

99. Schliehe C, Flynn EK, Vilagos B, Richson U, Swaminanthan S, Bosnjak B, et al. The methyltransferase Setdb2 mediates virus-induced susceptibility to bacterial superinfection. Nat Immunol. 2015;16(1):67–74. Epub 20141124. doi: 10.1038/ni.3046. PMID: 25419628.

100. Hansell CAH, Fraser AR, Hayes AJ, Pingen M, Burt CL, Lee KM, et al. The Atypical Chemokine Receptor Ackr2 Constrains NK Cell Migratory Activity and Promotes Metastasis. J Immunol. 2018;201(8):2510–9. Epub 20180829. doi: 10.4049/jimmunol.1800131. PMID: 30158126.

101. Galicia G, Maes W, Verbinnen B, Kasran A, Bullens D, Arredouani M, et al. Haptoglobin deficiency facilitates the development of autoimmune inflammation. Eur J Immunol. 2009;39(12):3404–12. doi: 10.1002/eji.200939291. PMID: 19795414.

102. Wang Y, Kinzie E, Berger FG, Lim SK, Baumann H. Haptoglobin, an inflammation-inducible plasma protein. Redox Rep. 2001;6(6):379–85. doi: 10.1179/135100001101536580. PMID: 11865981.

103. Naser W, Maymand S, Dlugolenski D, Basheer F, Ward AC. The Role of Cytokine-Inducible SH2 Domain-Containing Protein (CISH) in the Regulation of Basal and Cytokine-Mediated Myelopoiesis. Int J Mol Sci. 2023;24(16). Epub 20230814. doi: 10.3390/ijms241612757. PMID: 37628937.

104. Paul SP, Taylor LS, Stansbury EK, McVicar DW. Myeloid specific human CD33 is an inhibitory receptor with differential ITIM function in recruiting the phosphatases SHP-1 and SHP-2. Blood. 2000;96(2):483–90. PMID: 10887109.

105. Nassar M, Tabib Y, Capucha T, Mizraji G, Nir T, Pevsner-Fischer M, et al. GAS6 is a key homeostatic immunological regulator of host-commensal interactions in the oral mucosa. Proc Natl Acad Sci U S A. 2017;114(3):E337–E46. Epub 20170103. doi: 10.1073/pnas.1614926114. PMID: 28049839.

106. Lou Y, Li PH, Liu XQ, Wang TX, Liu YL, Chen CC, et al. ITGAM-mediated macrophages contribute to basement membrane damage in diabetic nephropathy and atherosclerosis. BMC Nephrol. 2024;25(1):72. Epub 20240227. doi: 10.1186/s12882-024-03505-1. PMID: 38413872.

107. Zhang H, Coblentz C, Watanabe-Smith K, Means S, Means J, Maxson JE, et al. Gain-of-function mutations in granulocyte colony-stimulating factor receptor (CSF3R) reveal distinct mechanisms of CSF3R activation. J Biol Chem. 2018;293(19):7387–96. Epub 20180323. doi: 10.1074/jbc.RA118.002417. PMID: 29572350.

108. Sipprell SE, Johnson MB, Leach W, Suptela SR, Marriott I. Staphylococcus aureus Infection Induces the Production of the Neutrophil Chemoattractants CXCL1, CXCL2, CXCL3, CXCL5, CCL3, and CCL7 by Murine Osteoblasts. Infection and immunity. 2023;91(4):e0001423. Epub 20230307. doi: 10.1128/iai.00014-23. PMID: 36880752.

109. Li R, Xie J, Xu W, Zhang L, Lin H, Huang W. LPS-induced PTGS2 manipulates the inflammatory response through trophoblast invasion in preeclampsia via NF-kappaB pathway. Reprod Biol. 2022;22(4):100696. Epub 20221031. doi: 10.1016/j.repbio.2022.100696. PMID: 36327673.

110. Gowhari Shabgah A, Jadidi-Niaragh F, Mohammadi H, Ebrahimzadeh F, Oveisee M, Jahanara A, et al. The Role of Atypical Chemokine Receptor D6 (ACKR2) in Physiological and Pathological Conditions; Friend, Foe, or Both? Front Immunol. 2022;13:861931. Epub 20220523. doi: 10.3389/fimmu.2022.861931. PMID: 35677043.

111. Tang R, Tay SS, Sharbeen G, Herrmann D, Youkhana J, Timpson P, et al. Bystander Expression of Atypical Chemokine Receptor 2 Protects T Cells from Chemoattraction towards Cancer-Associated Fibroblasts. Eur J Immunol. 2025;55(2):e202451215. doi: 10.1002/eji.202451215. PMID: 39931761.

112. Karve TM, Rosen EM. B-cell translocation gene 2 (BTG2) stimulates cellular antioxidant defenses through the antioxidant transcription factor NFE2L2 in human mammary epithelial cells. J Biol Chem. 2012;287(37):31503–14. Epub 20120405. doi: 10.1074/jbc.M112.367433. PMID: 22493435.

113. Cordani M, Sanchez-Alvarez M, Strippoli R, Bazhin AV, Donadelli M. Sestrins at the Interface of ROS Control and Autophagy Regulation in Health and Disease. Oxid Med Cell Longev. 2019;2019:1283075. Epub 20190507. doi: 10.1155/2019/1283075. PMID: 31205582.

