## Supplementary material for "Distinct temporal patterns of liver immune responses to pathogenic and non-pathogenic *Entamoeba histolytica* clones": S3 Table: S3 Table. FC_alle 151.docx

**Table S3. Differential gene expression of 151 immune-related genes in liver tissue following infection with A1ⁿᵖ and B2ᵖ trophozoites.**

|  |  | 6 hpi | | | | 12 hpi | | | | 24 hpi | | | |
| --- | --- | --- | --- | --- | --- | --- | --- | --- | --- | --- | --- | --- | --- |
|  |  | A1^np^ vs. Ctrl | | B2^p^ vs. Ctrl | | A1^np^ vs. Ctrl | | B2^p^ vs. Ctrl | | A1^np^ vs. Ctrl | | B2^p^ vs. Ctrl | |
|  | NAME | FC | FDR | FC | FDR | FC | FDR | FC | FDR | FC | FDR | FC | FDR |
| 1 | Ackr2  B2^p^ vs A1^np^ | 1.86 | ns | 4.69 | 2.55E-07 | 1.61 | ns | 4.87  2.72 | 8.33E-08  0.015 | -1.17 | ns | 1.08 | ns |
| 2 | Acod1 | 14.00 | 4.41E-06 | 14.11 | 7.63E-05 | 28.72 | 1.02E-09 | 25.14 | 3.74E-07 | 16.82 | 3.11E-06 | 33.57 | 1.50E-07 |
| 3 | Apcs | 2.76 | 0.0064 | 2.48 | 0.0016 | 2.55 | 0.012 | 1.50 | ns | 5.08 | 6.88E-06 | 4.13 | 4.57E-07 |
| 4 | Arid5a  B2^p^ vs A1^np^ | 29.05 | 3.13E-18 | 17.14 | 3.93E-17 | 4.79 | 0.00037 | 19.89  3.95 | 4.03E-19  0.0006 | 1.61 | ns | 2.23 | ns |
| 5 | Atf3 | 11.41 | 1.24E-11 | 16.41 | 2.33E-20 | 22.74 | 3.90E-19 | 38.27 | 5.67E-35 | 2.11 | ns | 2.65 | 0.023 |
| 6 | Btg2  B2^p^ vs A1^np^ | 11.92 | 3.46E-25 | 15.77 | 2.43E-28 | 3.23 | 7.70E-06 | 21.62  6.19 | 2.15E-35  1.10E-10 | -1.18 | ns | 1.19 | ns |
| 7 | C5ar1 | 1.75 | ns | 2.43 | ns | 4.00 | 0.0015 | 4.36 | 0.0018 | 2.69 | ns | 3.64 | 0.029 |
| 8 | Carmil1  B2^p^ vs A1^np^ | 7.50 | 4.58E-20 | 10.04 | 3.30E-33 | 3.38 | 1.84E-07 | 11.38  3.02 | 3.64E-37  0.0002 | 1.18 | ns | -1.14 | ns |
| 9 | Cblb  B2^p^ vs A1^np^ | 3.05 | 0.0001 | 5.21 | 4.55E-13 | 5.10 | 1.12E-09 | 13.09  2.27 | 3.73E-32  0.02 | 1.70 | ns | 2.21 | 0.0075 |
| 10 | Ccl2 | 1.65 | ns | 1.19 | ns | 3.84 | 0.009 | 3.96 | 0.0063 | 3.46 | ns | 6.33 | 0.00059 |
| 11 | Ccl3 | 4.37 | ns | 4.25 | ns | 27.53 | 0.004 | 25.85 | 0.0043 | 11.16 | ns | 28.26 | 0.014 |
| 12 | CCl4 | 4.20 | ns | 3.13 | ns | 5.41 | 0.047 | 7.95 | 0.016 | 5.59 | ns | 8.70 | 0.039 |
| 13 | Ccl6 | 3.80 | 1.37E-05 | 5.48 | 4.21E-12 | 2.90 | 0.0009 | 4.89 | 1.09E-10 | 2.66 | 0.011 | 2.60 | 0.0014 |
| 14 | Ccl7 | 1.35 | ns | 5.20 | ns | 11.15 | 0.011 | 6.49 | 0.057 | 17.52 | 0.0069 | 16.14 | 0.0077 |
| 15 | Ccl24 | 1.00 | ns | -1.14 | ns | -1.19 | ns | 1.12 | ns | 3.06 | 0.046 | 2.88 | 0.019 |
| 16 | Ccr1 | 11.89 | 2.81E-09 | 13.16 | 1.84E-07 | 14.65 | 6.50E-11 | 28.28 | 1.4E-12 | 11.61 | 2.09E-08 | 19.53 | 4.00E-09 |
| 17 | Ccrl2 | 2.32 | 0.0311 | 2.54 | 0.011 | 2.63 | 0.0082 | 3.78 | 5.58E-05 | -1.08 | ns | 1.14 | ns |
| 18 | Cd14 | 8.63 | 4.78E-11 | 9.38 | 1.05E-08 | 8.58 | 5.55E-11 | 13.28 | 1.14E-11 | 6.71 | 5.48E-08 | 20.19 | 1.45E-14 |
| 19 | Cd163 | 1.34 | ns | 3.09 | 1.82E-08 | 2.71 | 0.00083 | 2.80 | 3.0E-07 | 1.39 | ns | 1.17 | ns |
| 20 | Cd28 | 15.30 | 1.91E-07 | 22.05 | 8.53E-11 | 13.76 | 6.66E-07 | 25.54 | 5.74E-12 | 1.44 | ns | 1.33 | ns |
| 21 | Cd300lf | 17.58 | 1.97E-10 | 18.27 | 6.38E-10 | 11.45 | 1.40E-07 | 18.90 | 2.82E-10 | 5.06 | 0.0056 | 7.34 | 0.00028 |
| 22 | Cd33 | 6.37 | 2.10E-06 | 6.14 | 3.17E-06 | 8.21 | 3.23E-08 | 11.18 | 6.03E-11 | 3.92 | 0.0042 | 4.82 | 0.00031 |
| 23 | Cd44 | 1.16 | ns | 1.24 | ns | 1.65 | ns | 1.31 | ns | 1.85 | ns | 2.35 | 0.0015 |
| 24 | Cd46 | 1.95 | ns | 3.12 | 3.40E-05 | 4.76 | 6.53E-08 | 5.33 | 56E-11 | 3.51 | 0.00012 | 3.83 | 1.90E-06 |
| 25 | Cd53 | 2.32 | 0.010 | 2.51 | 0.0086 | 1.99 | 0.04 | 4.30 | 3.1E-06 | 1.95 | ns | 2.10 | ns |
| 26 | Cd63 | 1.21 | ns | 1.51 | ns | 3.23 | 0.00039 | 2.48 | 0.0062 | 2.28 | ns | 4.25 | 1.0354E-05 |
| 27 | Cd93 | 2.85 | 1.51E-05 | 3.58 | 6.05E-11 | 3.48 | 1.1E-07 | 4.89 | 5.37E-17 | 1.85 | ns | 2.01 | 0.004 |
| 28 | Cdkn1a | 11.63 | 1.57E-09 | 19.66 | 2.04E-22 | 15.36 | 1.075E-11 | 27.02 | 1.36E-27 | -1.23 | ns | 1.10 | ns |
| 29 | Cebpd  B2^p^ vs A1^np^ | 16.16 | 1.03E-21 | 17.40 | 1.02E-29 | 8.99 | 2.12E-13 | 20.83  2.27 | 8.80E-34  0.01 | 3.14 | 0.0026 | 3.06 | 0.00033 |
| 30 | Chil3 | 1.94 | ns | 2.69 | ns | 3.95 | 0.01 | 1.97 | ns | 26.05 | 2.715E-12 | 15.72 | 2.83E-09 |
| 31 | Chi3l1 | 6.14 | 0.02 | 10.66 | 0.0006 | 17.66 | 3.06E-05 | 14.20 | 4.89E-05 | 24.25 | 1.01E-05 | 22.03 | 6.00E-06 |
| 32 | Cish | 2.80 | 0.053 | 1.667 | ns | 5.40 | 0.00026 | 1.21 | ns | 3.86 | 0.02 | 3.71 | 0.026 |
| 33 | Clec4d | 7.85 | 0.00023 | 7.03 | 0.001 | 12.48 | 2.06E-06 | 10.86 | 2.25E-05 | 8.42 | 0.00046 | 20.08 | 2.05E-07 |
| 34 | Clec4e | 27.14 | 1.24E-05 | 26.34 | 1.69E-05 | 47.85 | 1.17E-07 | 45.69 | 1.47E-07 | 30.66 | 2.11E-05 | 67.23 | 3.22E-08 |
| 35 | Csf2ra | 1.87 | ns | 2.57 | 0.028 | 2.21 | ns | 4.20 | 0.00014 | 1.54 | ns | 2.55 | ns |
| 36 | Csf2rb | 2.42 | 0.0078 | 2.80 | 0.0026 | 2.20 | 0.018 | 2.35 | 0.012 | 1.31 | ns | 1.85 | ns |
| 37 | Csf3r | 2.59 | 0.023 | 2.85 | 0.026 | 3.97 | 0.00024 | 4.39 | 0.0004 | 3.45 | 0.005 | 4.16 | 0.0034 |
| 38 | Cxcl1 | 25.63 | 2.44E-29 | 8.25 | 5.60E-08 | 9.54 | 4.0E-14 | 9.85 | 1.81E-09 | 5.94 | 3.87E-08 | 12.09 | 3.54E-10 |
| 39 | Cxcl2 | 14.95 | 2.30E-05 | 14.26 | 2.26E-05 | 43.38 | 3.97E-10 | 80.62 | 2.64E-14 | 13.39 | 0.00023 | 48.87 | 2.49E-10 |
| 40 | Cxcl3 | 39.69 | ns | 67.21 | ns | 173.42 | 0.012 | 154.25 | 0.016 | 75.84 | ns | 496.97 | 0.01 |
| 41 | Cxcl9 | 1.63 | ns | 1.22 | ns | 1.14 | ns | 1.32 | ns | -2.38 | 0.022 | -2.39 | 0.01 |
| 42 | Cxcl10 | -1.22 | ns | 3.50 | 0.015 | -1.08 | ns | -1.47 | ns | 1.81 | ns | 1.22 | ns |
| 43 | Cxcl13 | 1.38 | ns | 1.09 | ns | 3.18 | 0.021 | 1.88 | ns | 1.25 | ns | 3.08 | 0.027 |
| 44 | Cxcr2 | 10.10 | ns | 8.88 | 1.34E-05 | 8.87 | 7.0E-07 | 11.70 | 3.58E-07 | 6.21 | 0.00026 | 7.24 | 0.00037 |
| 45 | Cxcr4 | 3.15 | 0.046 | 4.48 | 0.0094 | 5.97 | 0.00031 | 6.78 | 0.00029 | 5.07 | 0.0053 | 4.39 | 0.03 |
| 46 | Defb1 | -1.26 | ns | 1.65 | ns | 3.55 | 0.0014 | 5.53 | 7.62E-07 | -1.40 | ns | 1.86 | ns |
| 47 | Dusp1 | 3.18 | 0.030 | 5.36 | 3.68E-05 | 3.56 | 0.012 | 8.24 | 4.76E-08 | 1.49 | ns | 1.38 | ns |
| 48 | Dusp8 | 21.47 | 3.26E-09 | 30.82 | 7.21E-14 | 13.07 | 1.58E-06 | 32.00 | 2.70E-14 | -1.08 | ns | 1.24 | ns |
| 49 | Fas | 2.24 | 0.0052 | 2.47 | 0.00012 | 1.09 | ns | 2.04 | 0.0028 | 1.52 | ns | 1.71 | ns |
| 50 | Fgl1 | 2.03 | ns | 2.27 | 0.013 | 2.76 | 0.016 | 1.90 | 0.05 | 6.46 | 3.79E-06 | 3.93 | 1.79E-05 |
| 51 | Fos | 4.52 | 0.0025 | 6.54 | 5.12E-05 | 16.33 | 1.44E-10 | 28.96 | 1.088E-15 | 5.78 | 0.0009 | 8.95 | 4.32E-06 |
| 52 | Gadd45a  B2^p^ vs A1^np^ | 4.71 | 0.0004 | 6.19 | 1.92E-09 | 1.10 | ns | 4.77  4.03 | 3.24E-07  9.9E-05 | -1.45 | ns | -1.46 | ns |
| 53 | Gadd45b  B2^p^ vs A1^np^ | 26.43 | 2.17E-16 | 51.18 | 1.60E-36 | 17.87 | 1.39E-12 | 55.68  3.02 | 2.68E-38  0.003 | 2.09 | ns | 3.29 | 0.0051 |
| 54 | Gadd45g  B2^p^ vs A1^np^ | 47.85 | 1.06E-30 | 61.11 | 2.24E-87 | 17.82 | 4.69E-17 | 57.92  3.04 | 3.86E-85  0.006 | 6.78 | 6.15E-07 | 2.93 | 2.27E-05 |
| 55 | Gas6 | 1.63 | ns | 1.50 | ns | 1.17 | ns | 1.25 | ns | 3.58 | 7.96E-08 | 3.53 | 1.97E-09 |
| 56 | Gdf15 | 9.20 | 1.30E-05 | 16.87 | 7.58E-15 | 4.93 | 0.0032 | 13.17 | 1.46E-12 | -1.93 | ns | -1.68 | ns |
| 57 | Hp | 4.14 | 6.06E-07 | 4.52 | 9.02E-07 | 3.26 | 5.91E-05 | 4.22 | 2.14E-06 | 5.05 | 4.02E-08 | 4.50 | 4.27E-06 |
| 58 | Hamp | 2.20 | 0.0094 | 2.14 | 0.019 | 1.81 | ns | 2.69 | 0.00084 | -5.02 | 6.91E-09 | -1.89 | ns |
| 59 | Hmox1 | 4.35 | 2.35E-06 | 5.81 | 2.53E-18 | 6.58 | 3.9E-10 | 7.62 | 1.49E-24 | -1.31 | ns | -1.03 | ns |
| 60 | Icam1 | 3.06 | 6.38E-07 | 1.92 | 0.01 | 2.18 | 0.0012 | 1.97 | 0.005 | 1.63 | ns | 2.64 | 0.00011 |
| 61 | Ier3 | 7.48 | 9.62E-06 | 6.22 | 0.00022 | 10.06 | 1.59E-07 | 11.97 | 6.55E-08 | 3.91 | 0.023 | 10.81 | 1.53E-06 |
| 62 | Ifi27l2b | 1.23 | ns | 1.48 | ns | 1.90 | ns | 1.60 | ns | 3.65 | 0.00016 | 3.40 | 0.00033 |
| 63 | Ifrd1  B2^p^ vs A1^np^ | 9.07 | 7.58E-18 | 9.15 | 8.82E-23 | 8.11 | 4.83E-16 | 19.09  2.21 | 9.47E-41  0.01 | 2.53 | 0.0064 | 3.56 | 5.64E-07 |
| 64 | Igfbp1  B2^p^ vs A1^np^ | 35.70 | 4.20E-16 | 50.02 | 2.00E-24 | 2.39 | ns | 43.39  16.95 | 7.39E-23  3.27E-15 | 1.25 | ns | -1.88 | ns |
| 65 | Il13ra1 | 2.90 | 1.30E-07 | 3.02 | 8.03E-09 | 1.98 | 0.0017 | 3.15 | 1.35E-09 | 1.66 | ns | 1.43 | ns |
| 66 | Il17ra | 4.43 | 9.92E-10 | 4.12 | 5.54E-08 | 1.36 | ns | 3.10 | 1.95E-05 | 1.17 | ns | 1.35 | ns |
| 67 | Il1a | 1.50 | ns | 1.22 | ns | 2.01 | 0.048 | 2.03 | 0.032 | 1.45 | ns | 2.37 | 0.026 |
| 68 | Il1b | 1.97 | ns | 2.64 | 0.044 | 8.25 | 6.50E-11 | 10.72 | 2.28E-09 | 4.83 | 1.42E-05 | 14.90 | 5.36E-11 |
| 69 | Il1r1 | 14.63 | 6.30E-29 | 16.17 | 5.81E-52 | 10.77 | 1.26E-22 | 20.00 | 1.64E-60 | 1.76 | ns | 2.25 | 0.00035 |
| 70 | Il1r2 | 92.60 | 2.39E-05 | 141.64 | 3.36E-06 | 186.09 | 5.37E-07 | 311.01 | 2.31E-08 | 31.86 | 0.0094 | 106.77 | 5.39E-05 |
| 71 | Il1rn | 15.52 | 4.59E-25 | 12.62 | 1.21E-13 | 9.86 | 3.13E-17 | 10.35 | 9.66E-12 | 5.14 | 4.09E-08 | 9.44 | 5.97E-10 |
| 72 | Il18bp | 1.17 | ns | 1.08 | ns | 1.46 | ns | 1.18 | ns | 2.07 | 0.029 | 1.68 | ns |
| 73 | Irak3 | 7.45 | 1.57E-08 | 6.60 | 2.20E-07 | 6.09 | 6.11E-07 | 7.36 | 2.31E-08 | 2.08 | ns | 3.46 | 0.0057 |
| 74 | Itgam | 1.94 | 0.047 | 1.73 | ns | 2.78 | 0.0005 | 2.35 | 0.01 | 3.92 | 3.79E-06 | 4.28 | 1.14E-05 |
| 75 | Lbp | 2.14 | 0.00083 | 2.22 | 0.00065 | 2.19 | 0.00048 | 2.85 | 1.70E-06 | 2.31 | 0.0007 | 2.43 | 0.00038 |
| 76 | Lcn2 | 74.59 | 6.16E-43 | 49.90 | 2.56E-29 | 82.38 | 4.84E-45 | 66.76 | 4.68E-34 | 61.23 | 1.29E-38 | 73.94 | 1.81E-34 |
| 77 | Lepr | 25.60 | 2.59E-30 | 30.14 | 2.13E-39 | 49.79 | 2.0E-44 | 64.15 | 3.20E-59 | 2.11 | ns | 4.45 | 4.57E-07 |
| 78 | Lilrb4a | 1.84 | ns | 2.51 | 0.055 | 2.29 | ns | 2.96 | 0.014 | 1.71 | ns | 2.70 | ns |
| 79 | Lilrb4b | 5.70 | 4.56E-06 | 5.14 | 6.71E-05 | 6.77 | 2.82E-07 | 11.50 | 1.22E-10 | 3.91 | 0.0028 | 7.53 | 1.25E-06 |
| 80 | Map3k6 | 9.82 | 2.51E-09 | 16.22 | 1.40E-22 | 19.77 | 6.44E-16 | 31.97 | 2.46E-35 | 1.68 | ns | 1.42 | ns |
| 81 | Mapkapk2 | 2.27 | 0.0018 | 2.82 | 10.00E-07 | 1.57 | ns | 3.43 | 1.54E-09 | -1.21 | ns | -1.079 | ns |
| 82 | Mmp8 | 25.69 | 2.60E-07 | 39.86 | 4.52E-09 | 39.22 | 2.64E-09 | 47.82 | 4.10E-10 | 7.56 | 0.016 | 14.30 | 0.00025 |
| 83 | Mmp9 | 1.59 | ns | 1.54 | ns | 3.68 | 0.0063 | 5.86 | 0.00034 | 4.88 | 0.0021 | 5.28 | 0.0038 |
| 84 | Mt1 | 90.06 | 4.10E-28 | 92.93 | 1.65E-30 | 61.22 | 2.35E-23 | 107.63 | 8.82E-33 | 12.12 | 5.00E-08 | 11.07 | 4.46E-08 |
| 85 | Mt2  B2^p^ vs A1^np^ | 224.63 | 1.79E-34 | 239.29 | 1.49E-36 | 77.86 | 2.90E-22 | 251.00  3.04 | 1.89E-37  0.03 | 16.75 | 1.08E-08 | 16.58 | 4.87E-09 |
| 86 | Mx1 | 1.02 | ns | 1.52 | ns | 1.56 | ns | 2.07 | 0.049 | 1.13 | ns | 1.25 | ns |
| 87 | Nfkbia | 2.57 | 0.0025 | 3.53 | 1.10E-06 | 1.95 | 0.044 | 3.44 | 1.34E-06 | 1.45 | ns | 1.55 | ns |
| 88 | Nfkbiz | 3.25 | 1.41E-07 | 3.00 | 7.47E-06 | 3.58 | 7.26E-09 | 5.85 | 1.027E-14 | 2.24 | 0.0032 | 3.12 | 1.19E-05 |
| 89 | Ngp | 16.40 | 0.030 | 14.74 | 0.037 | 19.02 | 0.016 | 63.19 | 0.0002 | 2.63 | ns | 5.47 | ns |
| 90 | Nlrc5 | 2.37 | 0.0019 | 2.50 | 5.50E-05 | 4.58 | 10.0E-10 | 4.23 | 7.55E-12 | 2.10 | 0.033 | 2.03 | 0.0097 |
| 91 | Nlrp12 | 4.44 | 4.04E-08 | 5.38 | 3.36E-10 | 3.38 | 1.47E-05 | 4.19 | 1.051E-07 | 1.04 | ns | 1.79 | ns |
| 92 | Orm1 | 2.75 | 0.00055 | 3.06 | 3.12E-05 | 2.37 | 0.0041 | 2.33 | 0.0018 | 4.39 | 3.19E-07 | 3.39 | 1.59E-05 |
| 93 | Osm | 9.14 | ns | 21.57 | 0.014 | 71.03 | 0.00026 | 93.27 | 4.38E-05 | 30.36 | 0.02 | 121.18 | 7.56E-05 |
| 94 | Osmr | 3.37 | 5.20E-07 | 3.57 | 2.11E-08 | 2.50 | 0.00034 | 2.08 | 0.0027 | 1.71 | ns | 2.25 | 0.0033 |
| 95 | Phf11c  B2^p^ vs A1^np^ | 2.93 | 0.0023 | 12.73  4.23 | 3.36E-23  3.56E-07 | 9.97 | 3.99E-14 | 52.61  4.85 | 4.02E-57  1.94E-06 | 2.05 | ns | 3.05 | 0.00040 |
| 96 | Pik3ap1  B2^p^ vs A1^np^ | 5.66 | 2.27E-13 | 5.66 | 1.11E-20 | 1.94 | 0.017 | 5.17  2.44 | 9.45E-19  0.02 | -1.13 | ns | -1.17 | ns |
| 97 | Pla2g7 | 2.04 | 0.037 | 2.11 | 0.016 | 2.30 | 0.0089 | 2.00 | 0.02 | 1.78 | ns | 1.97 | ns |
| 98 | Plaur | 2.19 | ns | 3.29 | 0.042 | 5.48 | 0.0023 | 6.43 | 0.00024 | 3.89 | ns | 6.62 | 0.00092 |
| 99 | Plk3  B2^p^ vs A1^np^ | 29.26 | 1.58E-35 | 25.79 | 2.56E-42 | 10.63 | 2.42E-17 | 38.48  3.37 | 1.16E-53  9.27E-06 | 4.20 | 7.0E-06 | 2.97 | 0.0002 |
| 100 | Prtn3 | -3.34 | ns | 1.29 | ns | 1.22 | ns | -1.15 | ns | 15.48 | 7.33E-05 | 8.13 | 0.045 |
| 101 | Ptafr | 1.34 | ns | -1.09 | ns | 2.03 | 0.083 | 1.17 | ns | 2.89 | 0.016 | 3.43 | 0.0036 |
| 102 | Ptgs2 | 3.73 | ns | 3.68 | ns | 31.19 | 1.01E-06 | 22.43 | 3.73E-05 | 14.71 | 0.0012 | 78.78 | 9.20E-09 |
| 103 | Reg3b | 24.30 | 0.032 | 10.85 | ns | 42.82 | 0.0068 | 1.21 | ns | 4.39 | ns | 3.87 | ns |
| 104 | Retnlg | 10.06 | 9.90E-08 | 11.14 | 5.00E-06 | 10.50 | 4.80E-08 | 19.59 | 3.54E-09 | 8.05 | 1.03E-05 | 9.13 | 0.00013 |
| 105 | Rgs16  B2^p^ vs A1^np^ | 22.54 | 5.77E-10 | 55.54 | 8.62E-30 | 4.80 | 0.0057 | 83.49  15.81 | 2.23E-36  3.27E-15 | 1.36 | ns | 1.23 | ns |
| 106 | Ripk2 | 1.92 | 0.038 | 2.53 | 0.0028 | 2.51 | 0.0011 | 3.47 | 1.33E-05 | 1.77 | ns | 2.10 | 0.059 |
| 107 | S100a4 | 1.75 | ns | 1.88 | ns | 1.60 | ns | 1.43 | ns | 5.52 | 0.00044 | 4.11 | 0.0059 |
| 108 | S100a6 | 1.88 | ns | 2.31 | 0.022 | 2.09 | 0.052 | 1.88 | ns | 2.53 | 0.035 | 2.95 | 0.0047 |
| 109 | S100a8 | 12.52 | 1.26E-13 | 15.40 | 1.90E-10 | 11.96 | 4.27E-13 | 14.48 | 3.71E-10 | 9.68 | 3.02E-10 | 14.32 | 3.66E-09 |
| 110 | S100a9 | 10.39 | 4.00E-17 | 11.94 | 1.32E-09 | 12.40 | 9.99E-20 | 16.12 | 3.78E-12 | 9.22 | 1.43E-14 | 15.43 | 8.54E-11 |
| 111 | Saa1 | 25.82 | 3.46E-25 | 27.38 | 4.04E-35 | 18.36 | 5.22E-20 | 15.02 | 1.03E-23 | 53.86 | 1.79E-37 | 24.64 | 4.88E-32 |
| 112 | Saa2 | 72.42 | 5.75E-30 | 67.11 | 8.95E-50 | 61.12 | 1.30E-27 | 35.81 | 2.90E-36 | 113.96 | 1.40E-36 | 52.76 | 3.42E-43 |
| 113 | Saa3 | 6.71 | 4.88E-09 | 11.60 | 5.37E-08 | 12.18 | 1.79E-15 | 4.50 | 0.0016 | 16.46 | 9.65E-19 | 35.95 | 1.05E-15 |
| 114 | Scara5 | 7.23 | 5.00E-09 | 6.14 | 5.76E-08 | 3.057 | 0.003 | 4.62 | 5.56E-06 | 1.13 | ns | 2.29 | ns |
| 115 | Serpine1 | 101.31 | 1.07E-23 | 54.09 | 2.81E-16 | 158.89 | 2.38E-28 | 116.83 | 2.64E-23 | 29.06 | 9.86E-12 | 78.42 | 2.90E-18 |
| 116 | Serpina3c | 21.33 | 2.60E-09 | 30.85 | 2.89E-12 | 9.96 | 2.27E-05 | 11.43 | 1.92E-06 | 1.15 | ns | 2.59 | ns |
| 117 | Serpina3i | 52.56 | 3.53E-11 | 66.07 | 3.99E-11 | 23.32 | 4.03E-07 | 53.23 | 3.51E-10 | 5.69 | 0.056 | 22.65 | 1.09E-05 |
| 118 | Serpina3g | 6.05 | 4.68E-09 | 5.59 | 4.57E-10 | 2.42 | 0.012 | 2.63 | 0.001 | 1.04 | ns | 1.21 | ns |
| 119 | Serpina3n | 14.4 | 2.29E-23 | 13.58 | 4.07E-28 | 14.60 | 2.30E-23 | 12.63 | 1.01E-26 | 7.04 | 1.08E-11 | 8.53 | 6.24E-18 |
| 120 | Serpina3m | 8.17 | 2.47E-16 | 8.21 | 1.94E-18 | 6.73 | 2.18E-13 | 6.41 | 1.29E-14 | 3.38 | 4.53E-05 | 3.98 | 1.84E-07 |
| 121 | Serpinb2 | 3.90 | 0.052 | 1.61 | ns | 3.10 | ns | 2.11 | ns | 13.25 | 2.52E-05 | 22.47 | 9.46E-07 |
| 122 | Sesn1  B2^p^ vs A1^np^ | 1.49 | ns | 3.13 | 1.44E-07 | 1.44 | ns | 4.86  3.10 | 2.37E-14  0.001 | 1.23 | ns | -1.00 | ns |
| 123 | Setdb2  B2^p^ vs A1^np^ | 3.49 | 0.0001 | 12.35  3.46 | 4.92E-23  0.0003 | 5.39 | 2.98E-08 | 37.49  6.45 | 3.88E-48  3.84E-07 | 1.48 | ns | 1.70 | ns |
| 124 | Sftpa1  B2^p^ vs A1^np^ | 12.01 | 7.69E-07 | 14.33 | 1.47E-08 | 6.08 | 0.00085 | 18.27  2.80 | 2.70E-10  0.03 | 5.33 | 0.0094 | 5.77 | 0.0021 |
| 125 | Slfn1 | 3.66 | 0.01 | 2.73 | 0.052 | 4.18 | 0.003 | 3.29 | 0.012 | 4.85 | 0.0031 | 5.20 | 0.00095 |
| 126 | Slfn4 | 7.49 | 1.65E-05 | 8.01 | 2.61E-06 | 19.64 | 1.03E-11 | 10.58 | 3.63E-08 | 22.23 | 6.26E-12 | 24.73 | 7.50E-14 |
| 127 | Socs2  B2^p^ vs A1^np^ | 6.11 | 8.15E-10 | 4.87 | 1.55E-07 | 1.48 | ns | 3.65  2.28 | 2.11E-05  0.035 | -1.52 | ns | 1.48 | ns |
| 128 | Socs3 | 11.37 | 1.84E-18 | 9.90 | 1.88E-15 | 4.40 | 6.0E-07 | 8.35 | 1.95E-13 | 4.35 | 3.76E-06 | 6.24 | 3.49E-09 |
| 129 | Spp1 | -1.01 | ns | 1.62 | ns | 2.90 | 1.70E-06 | 2.97 | 3.30E-05 | 8.70 | 6.48E-26 | 8.26 | 9.57E-17 |
| 130 | Srgn | 3.92 | 2.42E-06 | 3.81 | 7.96E-06 | 4.76 | 3.90E-08 | 6.51 | 3.41E-11 | 3.70 | 3.58E-05 | 4.78 | 3.95E-07 |
| 131 | Stat3 | 2.99 | 3.11E-07 | 3.21 | 1.13E-07 | 1.89 | 0.0073 | 3.45 | 8.92E-09 | 1.44 | ns | 1.59 | ns |
| 132 | Steap4  B2^p^ vs A1^np^ | 18.62 | 3.94E-23 | 15.44 | 5.55E-16 | 2.38 | 0.015 | 9.79  3.82 | 2.22E-11  0.001 | 2.06 | ns | 3.50 | 0.0034 |
| 133 | Tab3 | 1.21 | ns | 1.52 | ns | 2.11 | 0.001 | 2.96 | 1.57E-07 | 1.42 | ns | 1.46 | ns |
| 134 | Tifa | 12.43 | 1.35E-25 | 8.31 | 1.23E-12 | 1.9 | 0.02 | 3.86 | 1.38E-05 | 1.97 | ns | 2.57 | 0.0169 |
| 135 | Tfrc | -1.79 | 0.0415 | -1.62 | 0.027 | -1.05 | ns | 1.04 | ns | 2.53 | 0.001 | 2.82 | 1.40E-07 |
| 136 | Tgfbr1 | 1.94 | 0.0019 | 1.69 | 0.0143 | 2.12 | 0.00026 | 2.14 | 7.74E-05 | 1.71 | 0.056 | 1.96 | 0.003 |
| 137 | Tgfbr2 | 2.04 | 0.0067 | 2.53 | 2.64E-06 | 2.49 | 0.00022 | 4.02 | 9.05E-14 | 1.01 | ns | -1.02 | ns |
| 138 | Thbs1 | 4.76 | 1.07E-06 | 4.98 | 1.19E-05 | 15.96 | 1.49E-20 | 13.24 | 3.4386E-14 | 6.87 | 3.24E-09 | 20.46 | 5.71E-18 |
| 139 | Timp1 | 10.00 | 0.00077 | 14.05 | 9.90E-05 | 8.58 | 0.002 | 11.34 | 0.00033 | 18.90 | 1.97E-05 | 32.16 | 3.018E-07 |
| 140 | Tlr2 | 4.05 | 2.42E-05 | 2.43 | 0.03 | 2.63 | 0.0068 | 2.44 | 0.022 | 1.85 | ns | 4.13 | 0.00039 |
| 141 | Tlr5 | 2.84 | 0.001 | 3.24 | 0.0001 | 4.19 | 1.39E-06 | 3.87 | 3.04E-06 | 1.91 | ns | 1.73 | ns |
| 142 | Tlr13 | 3.20 | 5.02E-05 | 2.70 | 0.00018 | 2.99 | 0.00016 | 3.17 | 5.54E-06 | 2.24 | 0.033 | 1.79 | ns |
| 143 | Tnfaip3 | 1.88 | ns | 1.57 | ns | 4.01 | 9.93E-06 | 3.45 | 0.0002 | 1.86 | ns | 3.24 | 0.0024 |
| 144 | Tnfrsf1a | 2.61 | 0.0001 | 2.51 | 0.00056 | 1.65 | ns | 2.04 | 0.008 | 1.21 | ns | 1.31 | ns |
| 145 | Tnfrsf1b  B2^p^ vs A1^np^ | 2.86 | 8.29E-05 | 3.63 | 1.37E-06 | 1.49 | ns | 4.17  2.58 | 3.85E-08  0.002 | 1.40 | ns | 1.37 | ns |
| 146 | Tnfrsf12a | 2.13 | ns | 2.90 | 0.00011 | 4.95 | 1.09E-05 | 4.25 | 2.08E-08 | 1.48 | ns | 2.03 | 0.051 |
| 147 | Traf6 | 1.676 | ns | 1.76 | 0.021 | 1.75 | 0.043 | 2.70 | 3.55E-06 | 1.31 | ns | 1.34 | ns |
| 148 | Trem1 | 28.04 | 9.47E-05 | 37.60 | 2.45E-05 | 71.12 | 1.28E-07 | 102.92 | 1.16E-08 | 40.38 | 3.74E-05 | 95.48 | 1.50E-07 |
| 149 | Txnip  B2^p^ vs A1^np^ | 12.13 | 2.04E-16 | 31.85 | 1.12E-47 | 7.85 | 3.79E-11 | 34.53  4.08 | 3.57E-50  0.001 | 1.00 | ns | -1.13 | ns |
| 150 | Vcam1 | 4.31 | 1.18E-10 | 5.16 | 2.29E-16 | 3.75 | 8.44E-09 | 4.24 | 6.72E-13 | 1.57 | ns | 1.47 | ms |
| 151 | Vnn1 | 2.12 | ns | 3.17 | 0.00057 | 2.31 | 0.041 | 3.12 | 0.00048 | 3.36 | 0.006 | 2.90 | 0.0056 |

Control: n = 5; infected groups: n = 4 per time point.

Gene expression changes are shown as fold change (FC) relative to uninfected control mice at 6, 12, and 24 hours post-infection (hpi).

Statistical significance was assessed using false discovery rate (FDR) correction.

Differential expression was defined as FC ≥ 2 and FDR < 0.05. Values not meeting these criteria are indicated as not significant (ns).

Red values indicate a significant difference between B2^p^ and A1^np^ infections at the respective time point.
